# Noradrenergic circuit control of non-REM sleep substates

**DOI:** 10.1101/2021.03.08.434399

**Authors:** Alejandro Osorio-Forero, Romain Cardis, Gil Vantomme, Aurélie Guillaume-Gentil, Georgia Katsioudi, Laura M.J. Fernandez, Anita Lüthi

## Abstract

One promising approach towards understanding what makes sleep vulnerable in disease is to look at how wake-promoting mechanisms affect healthy sleep. Wake-promoting neuronal activity is inhibited during non-REM sleep (NREMS). However, many mammalian species, including humans, show recurrent moments of sleep fragility during which sensory reactivity is elevated. Wake-related neuronal activity could thus remain active in NREMS, but its roles in dynamic variations of sensory reactivity remain unknown. Here, we demonstrate that mouse NREMS is a brain state with recurrent fluctuations of the wake-promoting neurotransmitter noradrenaline on the ∼50-seconds time-scale. These fluctuations occurred around mean noradrenaline levels greater than the ones of quiet wakefulness, while they declined steeply in REMS. They coincided with a clustering of sleep spindle rhythms in the forebrain and with heart rate variations. We addressed the origins of these fluctuations by using closed-loop optogenetic locus coeruleus (LC) activation or inhibition timed to moments of low and high spindle activity during NREMS. We could suppress, lock or entrain sleep spindle clustering or heart rate variations, demonstrating that both fore- and hindbrain-projecting LC neurons show synchronized infraslow activity variations in natural NREMS. Noradrenergic modulation of thalamic but not cortical circuits was required for sleep spindle clustering and involved noradrenaline release into primary sensory and reticular thalamic nuclei that activated both α1- and β-adrenergic receptors to cause slowly decaying membrane depolarizations. Noradrenergic signaling by LC, primarily known for attention promotion in wakefulness, renders mammalian NREMS more ‘wake-like’ on the close-to-minute-time scale through sustaining thalamocortical and autonomic sensory arousability.

## Introduction

The restorative and beneficial effects of sleep arise from its continuity.^1^ This requires that behavioral interactions with the sensory environment are suppressed. However, birds would crash and dolphins would drown if switching off from the sensory environment was not supplemented by vigilance.^2^ Since natural dangers and predators pose a risk to all animals irrespective of whether they are asleep or awake, it is natural to hypothesize that sleep evolution must have been tightly coupled to vigilance-promoting mechanisms. This may have placed natural healthy sleep dangerously close to vulnerability, such that minor shifts in the environment or in the organism’s physiology easily disrupt sleep. Indeed, neurological and psychiatric conditions underlying sleep disorders are highly diverse, yet identifying their origins remains challenging.^3–5^ For these reasons, a better estimate of sleep’s vulnerability in the healthy animal and of its neuronal basis is desirable.

During deep restorative non-rapid-eye-movement sleep (NREMS), the activity of wake-promoting areas ascending from subcortical areas into the forebrain is much reduced.^6^ However, already the first recordings of neuronal electrical activity in the sleeping animal indicated that not all these areas were entirely silent.^7, 8^ The locus coeruleus (LC), the most dorsal of ten noradrenergic nuclei located in the pontine brainstem,^7^ is strongly wake-promoting,^9^ tonically active during wakefulness,^7^ and becomes even more active in response to unexpected sensory stimuli.^10^ The LC also remains active during NREMS, although at minor rates that do not cause awakening.^7, 11^ This LC activity coincides with the appearance of EEG rhythms such as sleep spindles^7, 12, 13^ and cortical slow waves^11, 14^ and is relevant for sleep-dependent memory consolidation.^15^ Opto- or chemogenetic reinforcement of LC activity of rodents lowers auditory arousal thresholds in NREMS^16^ and increases functional connectivity in resting-state salience networks,^17^ which is a signature of enhanced vigilance. Therefore, natural variations of LC activity during NREMS could underlie NREMS vulnerability. To address this, we aimed to elucidate the real-time dynamics of LC activity and its functional impact on brain and bodily substates during NREMS. We used closed-loop optogenetic interference with LC activity in combination with local and global sleep recordings in mouse, fiber photometric assessment of norepinephrine (NE) levels, heart rate monitoring and the analysis of synaptic potentials generated by LC afferents *in vitro*. We demonstrate that noradrenergic activity in thalamic sensory nuclei is unexpectedly high in NREMS and fluctuates on an infraslow time scale. Moreover, LC coordinates central and autonomic physiological correlates that make NREMS alternate between substates of low and high arousability.

## Results

### Noradrenergic signaling during NREMS regulates the timing of sleep rhythms

Freely behaving mice sleep in NREM-REM sleep bouts interspersed by wakefulness during the light period (ZT0-12), which is their preferred resting phase. Figure 1A presents the sleep-wake behavior of a single mouse showing a hypnogram, obtained from polysomnography, and the corresponding time-frequency distribution derived from local field potential (LFP) recordings in the primary somatosensory cortex S1. We calculated power dynamics for two frequency bands characteristic for NREMS, the sigma (10 – 15 Hz) and the delta (1.5 – 4 Hz) frequency bands. As shown previously,^18^ prominent power fluctuations in the sigma but not the delta frequency band are present on an infraslow time interval with a peak around ∼0.02 Hz (∼50 s) (Figure 1B). The sigma frequency band is populated by sleep spindle rhythms^19^ that contribute to sensory decoupling during NREMS.^19, 20^ To determine whether sleep spindle density co-varied over 0.02 Hz, we used a previously developed spindle detection algorithm^21^ (Figure S1) and analyzed the phase-locking between sigma power and sleep spindle density in n = 33 mice (12 C57BL/6J and 21 dopamine-beta-hydroxylase (DBH)-Cre+/− mice) recorded during the light phase. Sleep spindles clustered at the peak of the 0.02 Hz-fluctuation in sigma power, whereas they were rare in the troughs (Figures 1C and 1D). Polar plots depicting the phase coupling of 168,097 spindles of 33 mice to the 0.02 Hz-fluctuation demonstrate the non-uniformity of this distribution (R of Rayleigh 0.51 ± 0.07; Figure 1D). Sleep spindles thus cluster on the 50-s time scale during NREMS, consistent with reports in mouse and human.^22, 23^ This indicates that NREMS fluctuates between spindle-rich and spindle-poor substates that are known to be related to different degrees of sensory arousability.^18^

**Figure 1.**
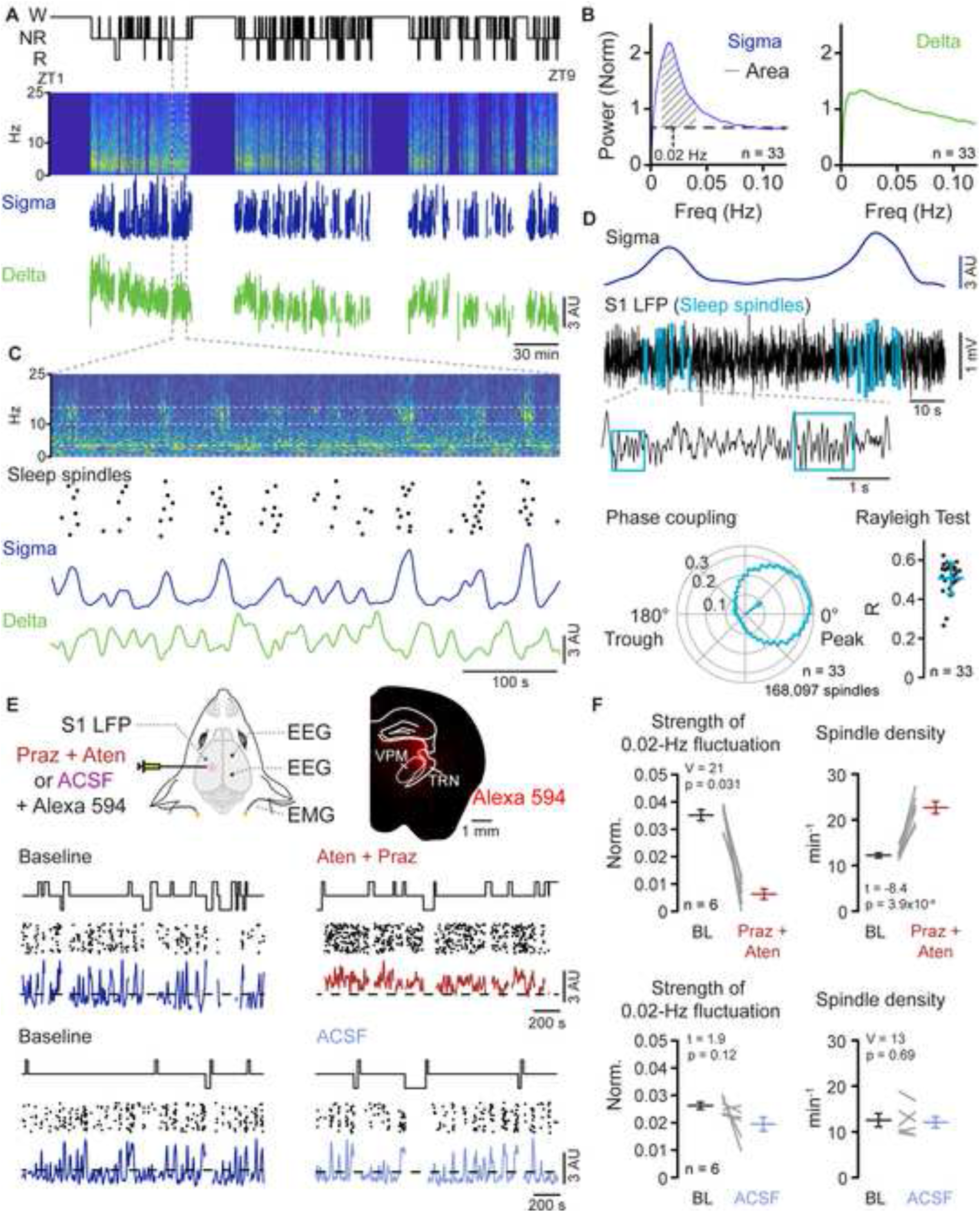
Noradrenergic signaling clusters sleep spindle rhythms during NREMS. (A) Hypnogram of a C57BL/6J mouse from ZT1-9, showing wake (W), NREMS (NR) and REM sleep (R), with corresponding time-frequency distribution of S1 LFP signal below. Power in NREMS only is indicated in color code, with warm colors indicating higher power. Summed sigma (10 – 15 Hz) and delta (1.5 – 4 Hz) power dynamics were derived from the time-frequency distributions. Dashed lines, NREMS bout selected for (C). (B) Fourier transform over sigma (left) and delta (right) power dynamics for n = 33 mice. Diagonal lines, area underneath the Fourier transform used to quantify the strength of the 0.02 Hz-fluctuation in this and subsequent Figure 2, Figure 3, and Figure 4. Vertical dashed line, 0.02 Hz. Horizontal dashed line, Mean values from 0.08-0.12 Hz. (C) Single NREMS bout indicated in (A). Dots, automatically detected spindle events. These are vertically jittered to visualize single spindles. (D) Example S1 LFP raw trace taken from (C) to show detailed position of spindles (see Figure S1). Note how sleep spindles (in blue squares) cluster when sigma power rises (top). ‘Phase coupling’: sleep spindle occurrence along the 0.02 Hz-fluctuation phases shown in a circular plot; arrow, mean Rayleigh vector across animals. ‘Rayleigh test’: quantification of the non-uniform distribution via R values across n = 33 mice (p < 1.0*10^−16^). (E) Experimental scheme for intracranial local pharmacology in combination with EEG/EMG and S1 LFP electrode implantation. Prazosin hydrochloride (Praz, 0.1 mM) and (S)-(-)-Atenolol (Aten, 5 mM) or artificial cerebrospinal fluid (ACSF) were co-injected with the red fluorescent dye Alexa594. Red labeling was verified post-hoc in sections from perfused brains. VPM, ventroposterial medial thalamus, TRN, thalamic reticular nucleus. Traces indicate hypnogram, sleep spindles and sigma power for two mice injected either with Aten + Praz (top) or ACSF (bottom). Dashed horizontal line, mean sigma power in Baseline. (F) Quantification of the strength of the 0.02 Hz-oscillation (as explained in (B)) and of sleep spindle densities, V, t and p values derived from Wilcoxon signed rank or paired t-tests, respectively.

To elucidate the role of noradrenergic signaling for infraslow variations in spindle density, we pharmacologically blocked noradrenergic receptors in thalamus, where sleep spindles originate.^19^ C57BL/6J mice were injected with a mix of α1- and β-noradrenergic antagonists (0.1 mM prazosin hydrochloride and 5 mM ((S)-(-)-atenolol, 150 nl) or control artificial cerebrospinal fluid (ACSF, 150 nl) locally into the somatosensory thalamus. Noradrenergic antagonist but not ACSF injections resulted in a rapid and reversible reduction of the strength of the 0.02-Hz fluctuation in sigma power in S1 (Figures 1E and 1F; Figure S2). Moreover, instead of being clustered, sleep spindles now appeared irregularly and at a mean density that was ∼2-fold higher (Figure 1F). The properties of individual spindle events including amplitude, intra-spindle frequency, number of cycles and duration, showed minor changes (Figure S3). Noradrenergic signaling in thalamus appears thus necessary for the generation of spindle-free periods and their repeated clustering on the infraslow time scale.

### Activity of the locus coeruleus is necessary and sufficient for spindle clustering during NREMS

To optogenetically interfere with the activity of the noradrenergic LC during NREMS in a time-controlled manner, we virally infected LC neurons of DBH-Cre mice to express excitatory (hChR2(H134R) (ChR2)) or inhibitory (Jaws) opsins. We then implanted these animals with EEG/EMG, an S1 LFP electrode and an optic fiber positioned uni- (for ChR2-expressing animals) or bilaterally (for Jaws-expressing animals) over the LC (Figures 2A and 2B). Optimal fiber positioning was ensured through intra-surgical pupil diameter monitoring (Figure S4). Using closed-loop monitoring of vigilance states, we first stimulated the LC specifically during NREMS at a low frequency (1 Hz) (Figure 2C) and confirmed the successful expression of the opsins post-mortem (Figure 2D). The 1-Hz frequency is within the range of spontaneous LC unit activity during NREMS^11, 12, 24^ and does not cause arousal in optogenetic studies.^9, 12^ Stimulation sessions took place in the first 20 min of each hour during 8 h of the light phase (ZT1-9), with light or sham (light source turned off) stimulation alternating over successive recording days. Light stimulation in NREMS produced a rapid onset and almost complete suppression of sigma power and of sleep spindles. The effect lasted as long as light was present and instantly recovered once optogenetic stimulation stopped (Figure 2E), decreasing the strength of the 0.02 Hz-fluctuation and sleep spindle density (Figure 2F). Conversely, continuous optogenetic inhibition according to the same experimental protocol locked sigma power at high levels and disrupted its fluctuation, increasing spindle density (Figures 2G and 2H). These results demonstrate that LC activity is both necessary and sufficient for the 0.02 Hz-fluctuation and the clustering of sleep spindles.

**Figure 2.**
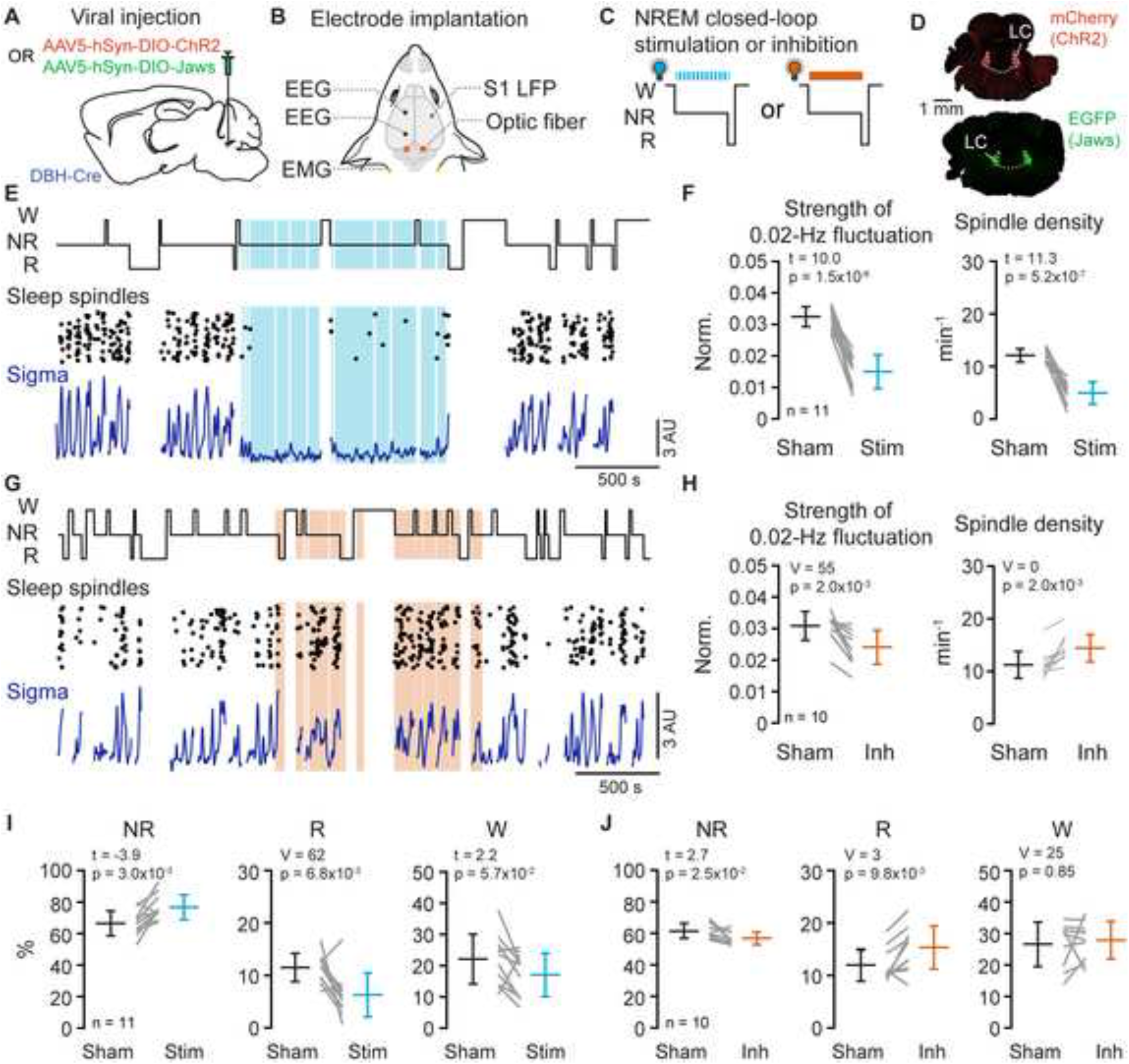
Optogenetic interrogation of the locus coeruleus during NREMS. (A) Viral injection strategy for DBH-Cre mice. For details of the viral plasmids, see Methods. (B) Experimental scheme for electrode implantation. (C) Schematic indicating closed-loop stimulation (blue) or inhibition (orange) of LC during NREMS (NR). LC optogenetic interrogation took place in the first 20 min of each hour during 8 h of the light phase (ZT1-9), with light or sham stimulation alternating over successive recording days. W, Wake; R, REMS. (D) Representative fluorescent microscopy images confirming viral expression in two mice included in the dataset. (E) Optogenetic LC stimulation (1 Hz, blue shading) applied during NREMS (NR), illustrating effects on sleep spindles and sigma power. (F) Quantification of effects on 0.02 Hz-fluctuation strength and on spindle density caused by LC stimulation during NREMS. Statistical evaluation via paired t-tests. (G and H) Same as (E,F) for optogenetic inhibition of LC. Statistical evaluation via Wilcoxon signed rank test. (I) Quantification of times spent in NR, R or W during LC stimulation or sham periods. (J) Quantification of times spent in NR, R or W during LC inhibition or sham periods. V, t and p values derived from Wilcoxon signed rank or paired t-tests.

To explore whether LC stimulation or inhibition affected sleep architecture, we quantified total time spent in the different vigilance states. We found that LC stimulation prolonged total time spent in NREMS at the expense of REMS and wakefulness (Figure 2I), whereas LC inhibition had opposite effects (Figure 2J). These architectural alterations were not accompanied by significant changes in relative delta power (stimulation: increase by 13%, p = 0.10; inhibition: decrease <1%, p = 0.90). Therefore, a low level of LC activity appears to consolidate NREMS. This could be explained by the incompatibility of monoaminergic signaling with REMS, which would cause NREMS to continue while we stimulated LC.^7, 13, 25^

### Locus coeruleus signaling in thalamus but not in cortex underlies sleep spindle clustering

From their thalamic site of origin, sleep spindle activity propagates to cortical circuits.^19^ As LC innervates both thalamic and cortical brain areas,^26^ we tested the involvement of both innervation sites in the effects observed by direct LC stimulation. We placed the optic fiber over somatosensory thalamus or S1. For S1, the optic fiber stub was glued to the S1 LFP electrode at a distance of 800 – 1,200 µm over the tip. Light stimulation of thalamic LC afferents reproduced the suppressive effects observed with direct LC stimulation (Figures 3A and 3B). In contrast, cortical stimulation was ineffective (Figures 3C and 3D). Thus, synaptic noradrenergic activity within the thalamus appears to sensitively control the clustering of sleep spindles measured in Figure S1, in agreement with the pharmacological results in Figure 1.

**Figure 3.**
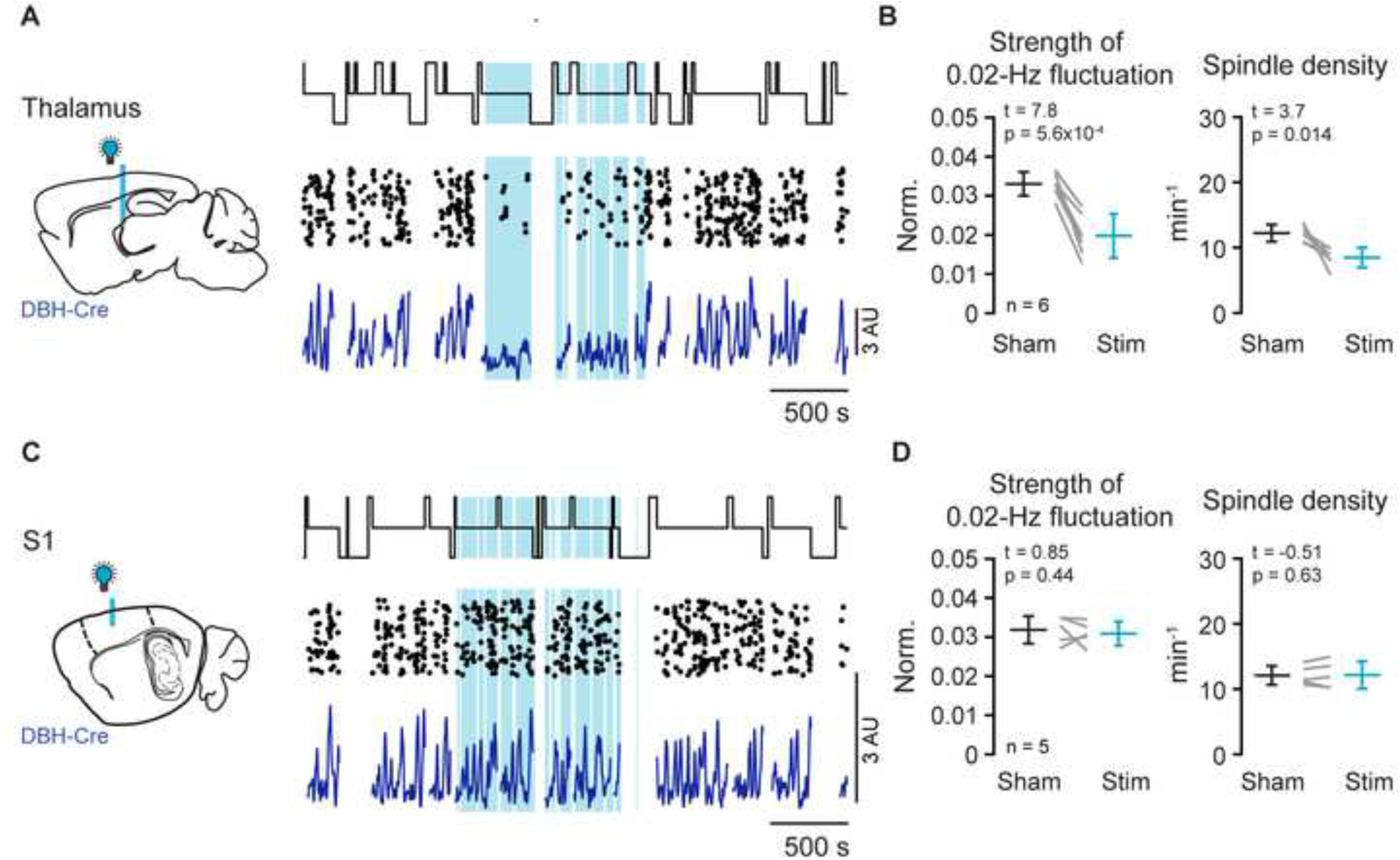
Optogenetic interrogation of locus coeruleus afferents in thalamus or cortex during NREMS. (A) Experimental scheme indicating optic fiber positioning over thalamus. For simplicity, EEG/EMG electrode and S1 LFP electrode (implanted as indicated in Figure 2B) are not shown. Representative traces on the right, similar arrangement of traces as in Figure 2E. (B) Quantification of effects on 0.02 Hz-fluctuation strength and on spindle density. (C and D) Same layout for experiments in which the optogenetic fiber was positioned over cortex. All p values in this Figure derived from paired t-tests.

### Locus coeruleus activity fluctuates on an infraslow time scale during NREMS

LC cells can discharge action potentials in both tonic and phasic modes during wakefulness, and both these modes have also been proposed to occur during NREMS.^7, 11, 13^ To address whether time variations in LC activity were relevant for the infraslow clustering of sleep spindles, we restricted the optogenetic manipulation of LC activity to distinct phases of the infraslow cycles. We detected these phases online through a machine-learning algorithm and triggered optogenetic activation based on whether sigma power started to rise or decline, thereby targeting preferentially high or low arousability periods^27^ (Figure 4). When we optogenetically activated LC whenever sigma power started rising, the 0.02-Hz fluctuation was suppressed (Figures 4A and 4B). This indicates that sigma increases, and sleep spindle generation, are not compatible with high LC activity. Conversely, when we inhibited LC when sigma power started declining, sigma power fluctuations became disrupted and tended to persist at high levels, thereby increasing sleep spindle density (Figures 4C and 4D). The decline in sigma power thus required LC activity. These two results are best compatible with LC activity increasing on infraslow time scales to suppress sleep spindles.

**Figure 4.**
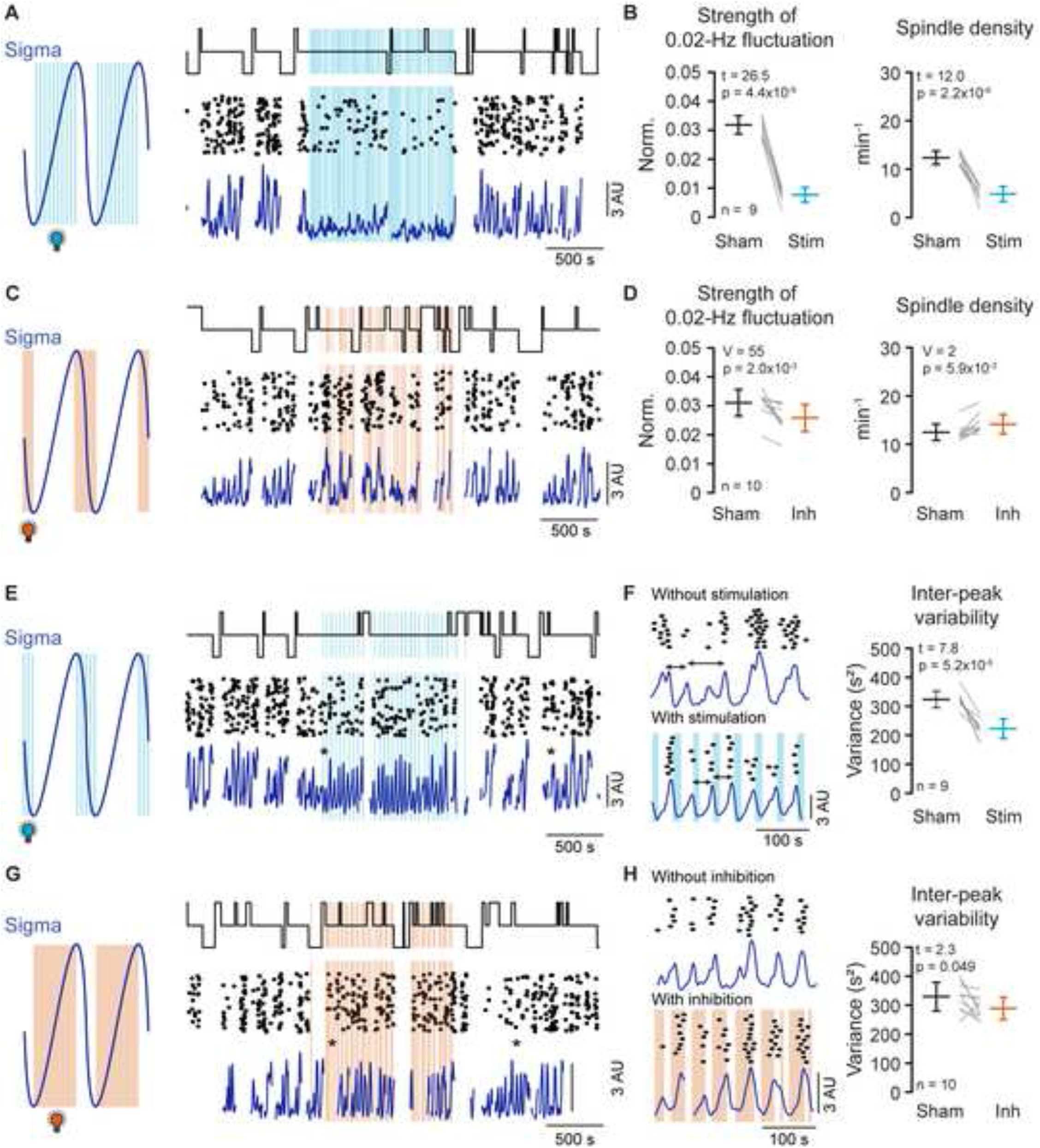
Optogenetic interrogation of locus coeruleus during spindle-rich or –poor periods of NREMS. (A) 1 Hz optogenetic stimulation of LC, restricted to moments of NREMS when sigma power increased. Blue waveform, schematic fluctuation to indicate the machine-learning-controlled timing of the light stimuli. Representative traces on the right, arranged as in Figure 2E. (B) Quantification of effects on 0.02 Hz-fluctuation strength and on spindle density. Statistical evaluation via paired t-tests. (C and D) Continuous optogenetic inhibition of LC, restricted to moments of NREMS when sigma power declined. Figure panels analogous to (A) and (B). V, t and p values derived from Wilcoxon signed rank test. (E) Optogenetic stimulation of LC when sigma power declined. Figure panels analogous to (A) and (C). (F) Expanded presentation of sigma and sleep spindle traces, taken from (*) in (E). Double-headed arrows denote interpeak intervals of the sigma power fluctuation. The regularization of the interpeak intervals in stimulation conditions is quantified through the variance (shown on the right). Note also the tighter temporal alignment between sleep spindles and sigma power. Statistical evaluation via paired t-tests. (G and H) Optogenetic inhibition of LC when sigma power increased. Figure panels analogous to (E) and (F). Statistical evaluation via paired t-tests.

Based on this result, we predicted that the converse experiment, stimulating LC when sigma power declined or inhibiting it when sigma power rose, would not disrupt infraslow dynamics. Intriguingly, for the first condition, successive cycles of high sigma power kept appearing (Figure 4E). Close inspection revealed that cycles appeared at shorter time intervals (sham: 52.9 ± 0.7 s, stim: 44.6 ± 1.7 s, n = 9, p = 5.0*10^−7^, paired *t*-test) and were more regular, as evident by the decreased peak-to-peak variability (Figure 4F). Strengthening LC activity when it was already naturally high thus entrained a regular and faster infraslow fluctuation. When we specifically inhibited LC activity during online detected periods of low sigma activity, an entrainment was again observed, with interpeak intervals shortened (sham: 53.7 ± 2.6, inhibition: 50.8 ± 1.5 s, n = 9; paired *t*-test, p = 3.6*10^−3^; Figure 4G) and regularized (Figure 4H). LC activity, already low at this moment, could thus be further inhibited by the light. This indicates that enforcing LC silence facilitates sigma level build-up and regularizes spindle clustering. Together, these results unravel a functionally relevant LC activity pattern during NREMS that interchanges between high, perhaps more phasic, and low, probably tonic, activity at infraslow timescales.

### Thalamic noradrenaline levels during NREMS are high and fluctuate on infraslow time intervals

LC activity is expected to release NE within thalamus and to stimulate noradrenergic receptors. However, the time course of free NE and of receptor signaling are unknown, leaving open how they determine the time course of sleep spindle dis- and re-appearance. We used fiber photometry to measure free NE levels in thalamic nuclei across the sleep-wake cycle by expressing the newly developed fluorescent NE biosensors GRAB_NE1h_ or GRAB_NE1m_ that have high and moderate affinity for NE, respectively.^28^ Mice expressing one of the two biosensors in thalamus were implanted for sleep monitoring and fiber photometry. The fluorescence signals varied across the three vigilance states, and showed higher mean values during NREMS compared to both, quiet wakefulness (QW) and REMS (Figures 5A and 5B). Vigilant-state-related changes showed a similar time course as measures in prefrontal cortex^13^ and are consistent with absent action potential activity in LC during REMS.^7^ In contrast, levels in NREMS were unexpectedly high. We observed rapid increases in NE levels when we stimulated the awake mouse in its cage by approaching one of our hands (Figure S5), indicating that we measured NE dynamics outside the saturating range of the biosensors.^28^ Focusing on NREMS bouts only, NE signals were inversely correlated with sigma power and showed recurrent negative peaks at infraslow intervals (Figure 5C). To resolve the time course of noradrenergic signaling on a cycle-to-cycle basis, we detected all infraslow sigma power cycles taking place during NREMS (excluding transitional periods) and examined the corresponding dynamics of free NE (Figure 5D). NE levels rose rapidly before sigma power declined, consistent with the suppressant effects of LC activity. Conversely, NE levels declined as sigma power was rising. Across animals, NE had already declined when sigma levels started rising, producing a non-symmetrical U-shaped time course. A similar profile of NE time course was observed for the medium-affinity biosensor GRAB_NE1m_ (Figure S5), indicating that the natural NE dynamics during NREMS were preserved across the range of affinities and on-/off-rates of the two sensors.^28^ Therefore, noradrenergic tone in thalamic sensory nuclei is higher than in quiet wakefulness and tightly regulates the dynamics of sleep spindles.

**Figure 5.**
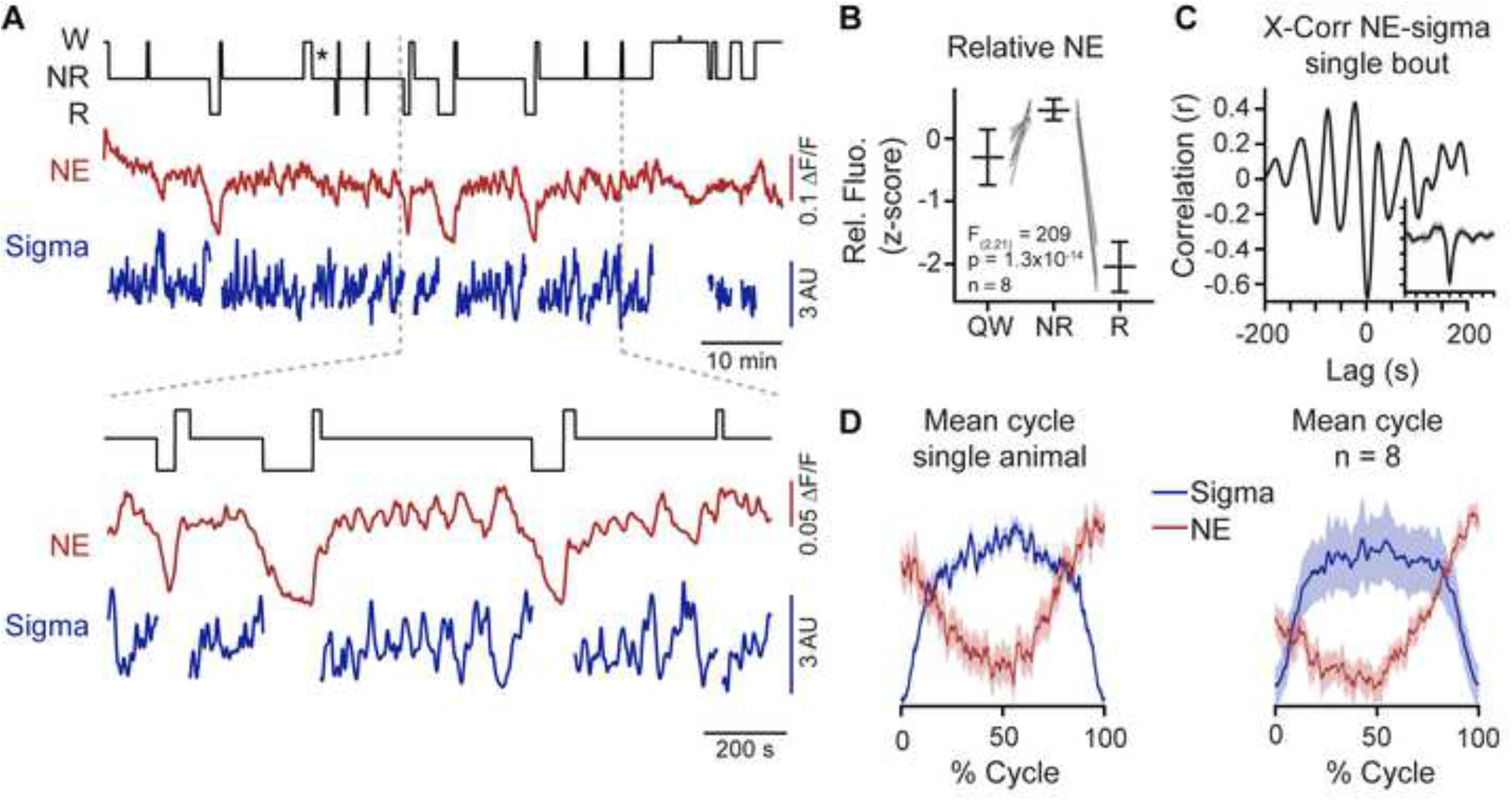
Free noradrenaline levels in thalamus are high and fluctuate during NREMS. (A) Representative recording showing (from top to bottom) hypnogram, relative fluorescence derived from the NE biosensor GRAB_NE1h_, and sigma power dynamics. Expanded portion shown below. *, bout selected for analysis in panel (C). Note the NE decrease during REMS and the higher NE levels during NREMS compared to quiet wakefulness. (B) Z-scored relative fluorescence derived from the NE biosensor GRAB_NE1h_ in three vigilance states quiet wakefulness (QW), NR and R for n = 8 animals. One-way ANOVA followed by post-hoc *t*-tests, which yielded: QW vs NR: t = −4.56, p = 2.6×10^−3^; QW vs R: t = 9.27, p = 3.35×10^−5^; NR vs R: t = 39.26, p = 1.81×10^−9^. (C) Cross-correlation (X-Corr) between sigma power and the NE biosensor signal for a single NREMS bout. Inset, Mean cross-correlation across all bouts in the recording, with axis scaling similar to the main figure of this panel. (D) Left, overlay of sigma power dynamics across all infraslow cycles detected from trough to trough, with corresponding NE fiber photometry signal for one mouse. Right, Mean traces across n = 8 animals.

### Ionic mechanisms underlying norepinephrine-induced membrane depolarizations

When exposed to NE, thalamic cells in vitro depolarize such that they no longer engage in spindle-like rhythms.^29^ However, the nature and the duration of synaptic communication between LC afferents and thalamic neurons are unknown. We hence combined patch-clamp recordings with optogenetic LC fiber stimulation and studied evoked responses in thalamocortical and thalamic reticular neurons, both of which are involved in sleep spindle generation.^19^ From DBH-Cre mice expressing ChR2 in LC, we prepared coronal thalamic slices and recorded from cells in the ventrobasal complex, which contains the somatosensory thalamus. Thalamocortical cells had resting membrane potentials between −65 to −70 mV, and responded with rebound burst discharge upon negative current injection (Figure S6). Optogenetic stimulation generated a slow membrane depolarization for stimulation frequencies at 1, 3 and 10 Hz (Figure 6A). Amplitudes of evoked potentials ranged between 0.8 – 4.5 mV with onset latencies of 1.25 – 6.9 s, and decayed with a slow time course lasting 66 – 106 s (Figure 6B, Figure S6). Only onset latency was modulated by stimulation frequency. The optogenetically evoked noradrenergic currents measured in cells voltage-clamped at −70 mV were blocked by atenolol (10 µM in bath), indicating involvement of β-adrenergic receptors (Figures 6C and 6D).^29^ Furthermore, the current response was largely eliminated by bath application of 1.5 – 3 mM Cs^+^ (Figures 6E and 6F), a blocker of cAMP-sensitive hyperpolarization-activated cation-channels.^30^ Both atenolol and Cs^+^ produced outward currents, indicating a standing receptor and current activation.

**Figure 6.**
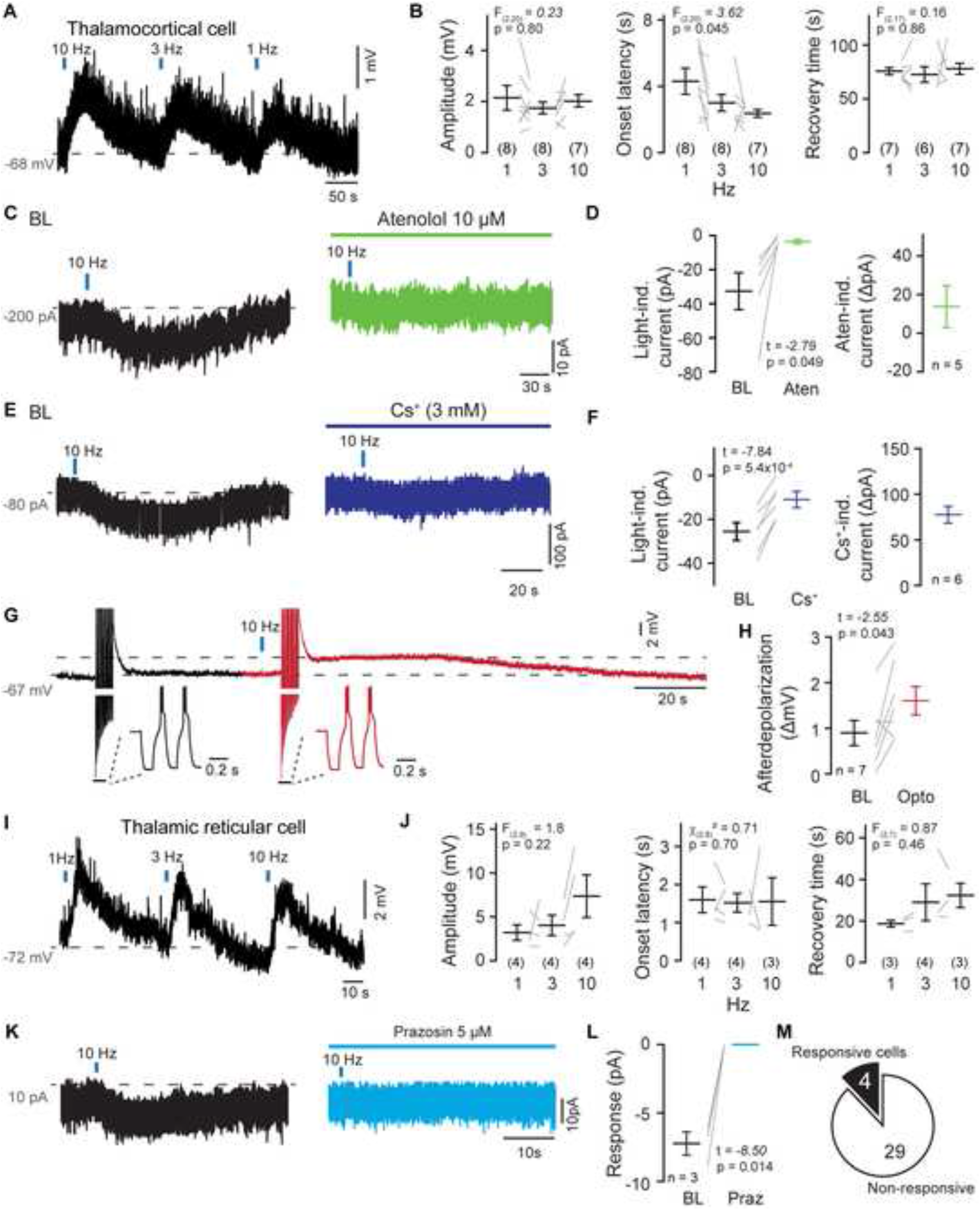
LC fiber stimulation-evoked slow noradrenergic potentials in thalamic neurons. (A) Representative recording from a whole-cell-patched thalamocortical cell exposed to three successive light stimuli at 10, 3 and 1 Hz (4 pulses, each lasting 100 μs). Dashed line, - 68 mV. (B) Quantification of evoked response amplitudes, onset latencies and recovery times as explained in (Figure S6). One-way ANOVA followed by post-hoc *t*-tests: 1 Hz vs 3 Hz: t = 1.6, p = 0.13; 1 Hz vs 10 Hz: t = 2.7, p = 0.024; 3 Hz vs 10 Hz: t = 1.1, p = 0.29. (C) Current response in a voltage-clamped thalamocortical neuron held at −70 mV to 10 Hz-light pulses before (left) and after (right) bath application of the β-adrenergic antagonist atenolol. Dashed line, −200 pA. (D) Left, Quantification of light-induced currents in baseline (BL) and Atenolol (Aten). Right, Aten-induced positive holding current shift. (E and F) Same as (C and D) for bath-application of Cs^+^ to block the cAMP-sensitive hyperpolarization-activated cation channels. Dashed line, −80 pA. (G) Example recording from a thalamocortical neuron injected with repetitive negative current pulses (−250 pA, 20 pulses, 120 ms each) to evoke low-threshold Ca^2+^ bursts. Insets show, each, two of these evoked bursts. Burst were followed by an afterdepolarization that was prolonged when light pulses (10 Hz, 4 pulses, blue bar) preceded current injections. Dashed lines aligned to baseline membrane potential and peak of the afterdepolarization. (H) Quantification of afterdepolarizations in baseline (BL, without light exposure) and with light exposure (Opto). (I and J) As (A) and (B), for a representative recording from a thalamic reticular neuron. Dashed line, −72 mV. F and χ^2^ derived from ANOVA and Kruskal-Wallis test, respectively. (K) Current response in a voltage-clamped thalamic reticular neuron held at −70 mV to 10 Hz light pulses before (left) and after (right) bath application of the α1-adrenergic antagonist prazosin. Dashed line, 10 pA. (L) Quantification of current response amplitude in baseline (BL) and during prazosin (Praz). (M) Number of TRN cells responding to LC optogenetic fiber stimulation. Statistical evaluation in (D, F, H and L) derived from paired t-tests.

To estimate the time course of action of NE on spindle-related cellular activity, we combined optogenetic stimulation of LC fibers with negative current injections to generate repetitive low-threshold burst discharges,^30^ known to occur during sleep spindles (Figure 6G).^19^ This resulted in a persistent afterdepolarization that was larger and longer than the one generated by cellular bursting alone (Figure 6H). The coincidence of sleep spindle activity with NE release thus generates a prolonged period of cellular depolarization, known to be sufficient to render thalamocortical cells refractory to synaptically driven burst discharge, which is necessary to engage in a next sleep spindle.^30^

Light-induced depolarizations were also observed in thalamic reticular cells recorded in the somatosensory sector of TRN (Figures 6I and 6J). Corresponding currents were largely blocked by the α1-adrenergic antagonist prazosin (5 µM in bath) (Figures 6K and 6L).^29^ However, < 15% of TRN cells showed a detectable current response (Figure 6M), suggesting that TRN cells could be heterogeneous in terms of noradrenergic responsiveness. Thalamic reticular cells thus also respond with slowly decaying membrane depolarizations when exposed to NE release from LC fibers.

### The locus coeruleus coordinates heart rate variations with infraslow brain rhythms

During mouse NREMS, the heart rate (HR) also fluctuates on an infraslow time scale and is anticorrelated to sigma power.^18^ We conjointly monitored HR and sigma power in freely sleeping C57BL/6J mice (Figures 7A and 7B) and used peripheral cardiac pharmacology to determine which branch of the autonomic nervous system controlled the infraslow variations in heart rate. The HR variations were suppressed by the peripheral parasympathetic antagonist methylatropine (10 mg kg^−1^) (Figures 7C and 7D)^31^ but not by the peripheral sympathetic antagonist atenolol (1 mg kg^−1^) (Figures 7E and 7F)^32^.

**Figure 7.**
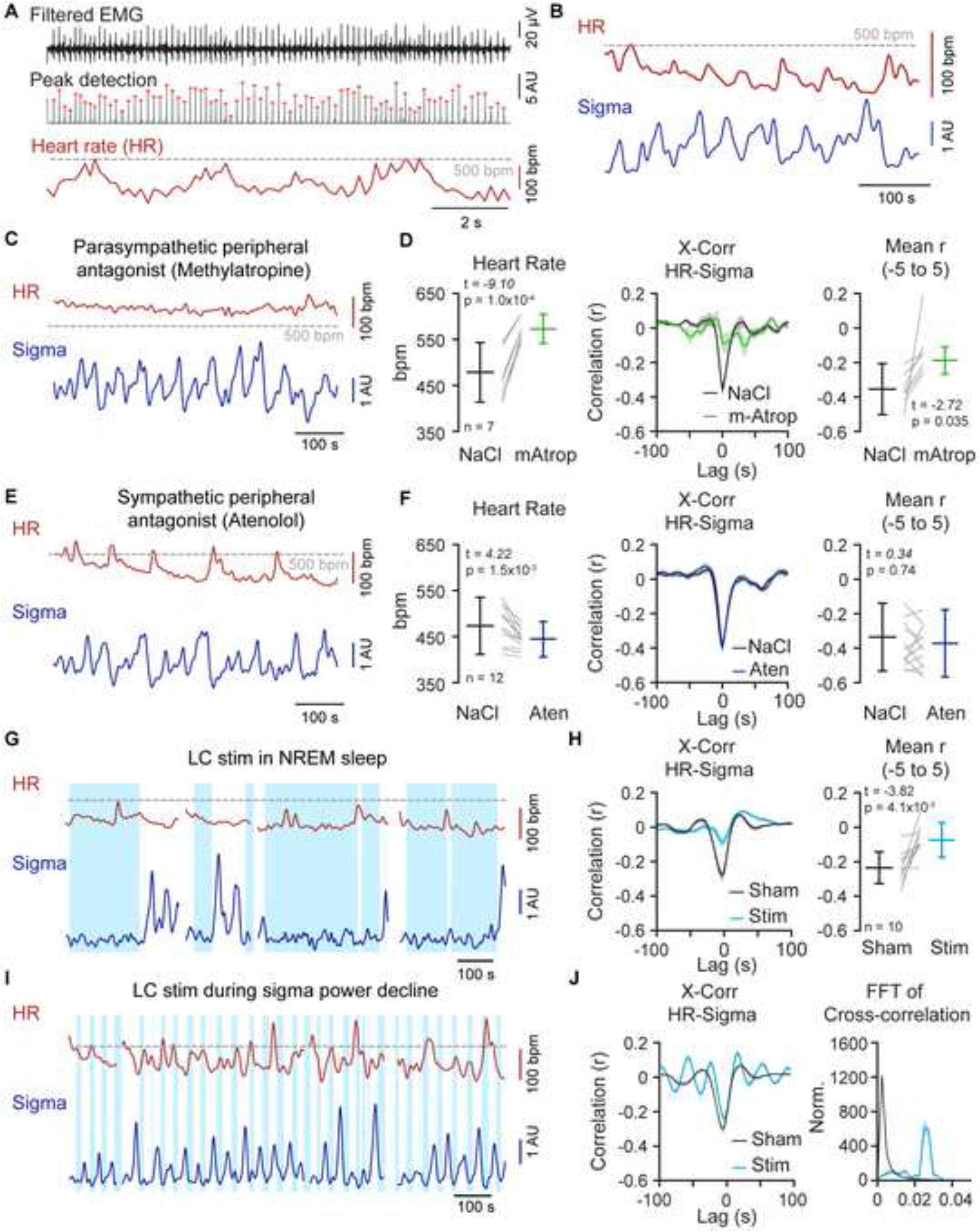
The LC coordinates heart rate and infraslow brain rhythms. (A) Extraction of HR from EMG traces. Top, raw high-pass-filtered (25 Hz) EMG, used for peak detection (middle) and HR (bottom) calculation. (B) Representative NREMS traces showing sigma power dynamics and corresponding HR after an i.p. injection of NaCl. (C) Example traces illustrating the effects of the parasympathetic antagonist methylatropine (mAtrop, 10 mg kg^−1^) on HR and sigma power dynamics. Dashed line, 500 bpm. (D) Left, Quantification of mean HR following NaCl or mAtrop injections; Middle, Corresponding cross-correlations (X-Corr), Right, Values of the cross-correlation between −5 to +5 s. (E and F) As (C) and (D) for injection of a sympathetic peripheral antagonist (Atenolol, 1 mg kg^−1^). (G) Example traces illustrating the effects of LC stimulation (stim) in NREMS on HR and sigma power dynamics. (H) Corresponding cross-correlations quantified as in (D) and (F). (I) As (G), for LC stimulation restricted to machine-learning-detected periods of declining sigma power. (J) Corresponding cross-correlation and its Fourier transform highlight the appearance of an infraslow peak. Statistical evaluation in this figure obtained from *t*-tests.

LC activity has been implied in parasympathetically driven HR variability in humans^33^ and, in rodents, augments inhibitory input to preganglionic cardiac vagal neurons.^34^ Therefore, we next tested whether optogenetic manipulation of LC affected variations in HR. Indeed, continuous LC stimulation during NREMS disrupted the infraslow HR variations (Figure 7G) and decreased their anticorrelation with sigma power (Figure 7H). To directly evaluate the capability of LC in entraining HR variations, we stimulated LC specifically when sigma power declined, as done before (see Figure 4E). This visibly augmented HR variations and imposed anticorrelations with sigma power with side-peaks showing an infraslow periodicity (Figures 7I and 7J). These data show that the LC is a source of HR variability during NREMS on an infraslow timescale. Moreover, LC is capable of coordinating sigma power and HR in a manner that supports a critical role in the generation of arousability variations during NREMS.

## Discussion

Noradrenergic cell groups in the pontine brainstem are conserved across fish, amphibians, reptiles, birds, and mammals.^35–37^ Across species, noradrenaline signaling plays universal roles during wakefulness, attention and stress, ensuring that important sensory stimuli are adequately detected and reacted to.^38^ Here we find that noradrenergic signaling during NREMS remains as high as in wakefulness in thalamus, the principal subcortical area implicated in sensory processing. Moreover, at intervals just under a minute (∼50 s), noradrenergic signaling drives NREMS between two substates previously associated with high and low sensory arousability.^18, 39^ Noradrenergic signaling is thus an integral part of mammalian NREMS, although numerically LC unit discharge occurs at much lower rates compared to wakefulness.^7, 24^ Our findings provide quantitative evidence for mammalian NREMS being associated to a high and dynamically varying noradrenergic tone, leveraging its sensory vigilance to levels found in wakefulness. Therefore, consistent with the original general hypothesis, mammalian NREMS has an innate and high vulnerability to disruption that arises from powerful wake-promoting circuit control of NREMS substates.

### The infraslow time scale is present in NREMS across species up to humans

The infraslow time scale has been repeatedly observed in diverse dynamic measures of mammalian NREMS^40^ and also appears in reptile sleep in the switch between two electrophysiologically distinct sleep states.^41^ The infraslow time interval could thus be a phylogenetically preserved temporal unit in sleep architecture that arises out of temporally fluctuating noradrenergic signaling. This possibility could be clarified in future comparative studies on species-specific noradrenergic signaling in sleep.^2^ In functional imaging studies of human deep NREMS (stage N3), the brainstem area surrounding the LC showed activity increases lasting ∼20 s when slow and delta waves were present.^14^ Both fast and slow spindles of human NREMS are nested to these waves in tight temporal sequence.^19^ Moreover, fast spindle activity is clustered on the infraslow time scale, in particular during light NREMS (stage N2).^18, 22, 42^ There is thus evidence for time scales compatible with recurrent LC activity increases over tens of seconds in humans. A recent study pointed out that the clustering of sleep spindles on similar time scales offered moments during which targeted memory reactivation was particularly effective when humans took a nap.^42^ This further supports the idea that variations in spindle activity divide NREMS into substates that differ in the way they engage in memory trace reactivation.^43^ Our study could also renew efforts to identify the unknown origins of the human-specific cyclic alternating patterns in the EEG. These events occur preferentially at phases of transitions between sleep stages and lead to higher arousability every 20 – 40 s.^44^ Intriguingly, the integrity of cyclic alternating patterns has been recently linked to the preservation of noradrenergic terminal density in LC and forebrain.^45^ Periodic leg movements that are widespread arousal events in the human sleep are yet another type of sleep-related arousal event on infraslow time scales.^46^ Currently available insights thus indicate that noradrenergic signaling is likely important for NREMS across species and in particular in humans, playing critical roles for NREMS substates and the control of arousability, including its alterations in neurodegenerative disorders.

### Infraslow noradrenergic signaling regulates cellular and circuit heterogeneity during NREMS

The discharges of LC units during NREMS presented in pioneering studies seem minor compared to wakefulness.^7, 11, 12^ We now find here robust pulsatile NE increases during NREMS on an infraslow time scale that are higher than NE levels in awake quiet mice. Free NE varies similarly in prefrontal cortex during mouse NREMS,^13^ suggesting that a fluctuating neuromodulatory tone of NE covers large portions of the forebrain. Supporting this, LC unit measures in anesthetized and sleeping rodents indicate that there is a considerable degree of synchronous discharge on the infraslow time scale. In rats under urethane anesthesia, the firing patterns measured from forebrain-projecting LC units exhibited infraslow variations that had stable phase relationships between unit pairs.^37, 47^ In freely sleeping mice, large peaks in fiber photometric measures of intracellular Ca^2+^ levels in LC clearly recurred about once a minute,^13^ suggesting that numerous LC units are synchronously active. Furthermore, we found stable phase relationships between forebrain spindle and heart rate infraslow fluctuations in NREMS. This would require LC units to become synchronized across dorsal and ventral regions of the LC, in which forebrain- and hindbrain-projecting cells preferentially reside, respectively.^48^ Although such scenarios remain to be assessed, our findings are in favor of a synchronous activity mode of LC during NREMS that imposes fluctuating levels of NE across both forebrain and hindbrain.^37^

Mild chemogenetic excitation of major portions of noradrenergic LC neurons in lightly anesthetized mice increased resting-state functional network connectivity measured using functional magnetic resonance imaging, in particular in the salience and dorsal attentional networks.^17^ Infraslow NE signaling during NREMS could thus also lead to activation of these hemodynamically defined functional circuits.^40^ It will be important to test whether the enhanced arousability observed during NREMS with low spindle activity,^18^ in which natural LC activity is high, is accompanied by enhanced functional connectivity in large-scale resting-state networks.

At the cellular level, slow waves and sleep spindles during NREMS regulate intracellular Ca^2+^ activity of cortical excitatory pyramidal neurons and interneuron somata^49^ and dendrites^50^ in a layer-specific manner, whereby sleep spindles play important roles in boosting activity in subgroups of pyramidal cells.^51^ Fluctuations in NE levels during NREMS could thus be critical in setting the time windows for cortical operations such as memory consolidation. Besides LC, other neuromodulatory systems are known to be active during NREMS.^8, 52^ An interesting case is the coincidence of noradrenergic and cholinergic activity that occurs for tens of seconds prior to NREM-REMS transitions^6, 53, 54^ and that gives rise to brief moments of high spindle activity, while hippocampus starts to generate REMS-related theta rhythmicity.^55^ These moments of coincidence between NREMS- and REMS-related activities appear to be critical for declarative memory formation.^19^

### LC-dependent control of sensory arousability during NREMS

LC activity likely controls sensory arousability through several mechanisms. First, LC responds to sensory stimuli,^56^ therefore, if LC neurons depolarize and discharge more action potentials, sensory throughput will be facilitated, as demonstrated recently using optogenetic LC stimulation.^16^ Second, LC-mediated sleep spindle suppression removes inhibitory constraints on the successful propagation of sensory inputs across the thalamocortical axis.^19^ Third, sensory-evoked discharge of single or multiple thalamic and cortical units is strengthened by LC stimulation, with thalamic neurons increasing sensory responsiveness more robustly (reviewed in^10^). The density of LC fiber varicosities is higher in somatosensory thalamus compared to cortical areas in rat^26^ and DBH immunoreactivity is overall highest in sensorimotor over other cortical areas in human.^57^ Noradrenergic signaling reconfigures neuronal activity within large-scale brain networks to sharpen vigilant attention during wakefulness in response to unexpected stimuli or acute stress.^17, 58^ These findings offer several entry points to further examine how noradrenergic modulation of thalamic circuits facilitates the processing of sensory stimuli during states of NREMS.

### Noradrenergic signaling does not terminate spindles but regulates their clustering

Sleep spindles were initially observed to coincide with LC activity during NREMS. Based on sleep spindles measured in EEG^7^ and in hippocampal LFPs,^12^ it has been repeatedly proposed that the LC terminates sleep spindles. We establish here a direct mechanistic opposition between NE levels and the circuits in which sleep spindles originate and can refine this widespread interpretation. First, we rapidly and reversibly suppressed local sleep spindles^21^ via optogenetic LC activation and second, we find that free NE levels ran opposite to locally generated sleep spindles. Third, we demonstrate that LC fiber activation promoted membrane depolarization consistent with thalamic circuit refractoriness.^59^ Furthermore, we could not observe major alterations in sleep spindle properties when we pharmacologically antagonized noradrenergic signaling in thalamus, although our spindle detection algorithm reliably discriminated between cortical area-specific sleep spindle properties.^21^ Furthermore, synaptic events caused by NE release rose slowly over seconds and are thus unlikely to shortcut individual spindle-events. Based on these combined cellular and in vivo data, we propose that noradrenergic signaling is tailored to generate prolonged relatively spindle-free periods through depolarizing thalamic neurons,^30^ but it does not act rapidly enough to terminate individual spindles.

### The cellular and ionic mechanisms underlying noradrenergic control of sleep spindle clustering

Although cellular and glial mechanisms with infraslow dynamics have been observed,^30, 60^ causal links to infraslow dynamics in vivo have not been made. Through combining in vivo and in vitro approaches, we identify an infraslow time course of action for NE released from LC terminals that likely underlies sleep spindle variations. NE induces a slowly decaying membrane depolarization through activation of both α1- or β-receptors in thalamocortical and thalamic reticular neurons, which retards the re-engagement of these cells in sleep spindle generation even after free NE levels have returned to baseline. In further support of this interpretation, we could entrain rhythmically the infraslow fluctuations when we reinforced or attenuated LC activity at appropriate moments. The reinforcement of LC activity most likely triggered membrane depolarizations more consequentially across large cell populations. Conversely, the attenuation of LC activity likely removed spurious LC activity during the continuity periods and allowed a more synchronous entry and exit of thalamic circuits within the infraslow cycles. While LC activity thus regulates spindle clustering at the population level, other cellular and synaptic mechanisms could be more involved in the termination of spindle-activity in individual cells (reviewed in ref.^19^). Furthermore, we cannot exclude that the optogenetic interference with LC activity modified cortical and/or brainstem feedback afferents onto LC^10^ that could have accelerated its infraslow activity.

To the best of our knowledge, the slow LC-triggered membrane depolarizations described here in the thalamus are the sole postsynaptic effects mediated exclusively by NE described so far. LC afferents to parabrachial nucleus release glutamate to generate fast glutamatergic currents,^61^ and the release of NE remained minor unless fibers were stimulated at 20 Hz. Thalamic LC-fiber-elicited responses were present and relatively uniform in amplitude and time course over the 1 – 10 Hz frequency range. The stability of the response across stimulation frequencies will ensure that thalamic membrane potentials are homogeneously depolarized across many neurons. Ultrastructural studies indeed suggest that noradrenergic terminals in the rodent ventrobasal thalamic complex do not form well-defined synaptic contacts,^62^ suggesting that released NE may diffuse from the site of release into the extracellular medium. Supporting this, we find that noradrenergic receptors activated by optogenetic LC fiber stimulation are the same as the ones targeted by bath-applied NE.^29^ There was also a measurable noradrenergic antagonist-sensitive holding current component in our slices, consistent with an ambient NE level generating a tonic noradrenergic signal.

### Noradrenergic signaling is a source of heart rate variations during NREMS

Initial studies pointed out that an infraslow periodicity was also present in the HR^18^ and in pupil diameter during NREMS.^39^ At moments when spindle density was low and arousals more likely, HR was high and pupil diameter large. Using peripherally acting drugs, we identified the parasympathetic system as key for the infraslow variations in HR, as it was also observed for pupil diameter variations.^39^ The LC increases HR through several pathways, amongst which there is a suppression of activity in the parasympathetic preganglionic vagal nuclei,^34^ which regulates HR variability in humans.^33^ Our study is the first to directly probe the consequences of specific optogenetic LC stimulation on HR, and these need to be further explored regarding frequency dependence, mechanisms, and variation with vigilance state. Continuous 1 Hz-stimulation during NREMS abolished infraslow HR variations, demonstrating that LC coordinates the infraslow activity patterns in brain and heart. Appropriate timing of LC stimulation also increased the fluctuations of the HR and strengthened anticorrelations on the infraslow time scale. We thus establish the LC as a coordinator of brain and bodily fluctuations during NREMS, which likely involve parasympathetic signaling. The neural systems underlying this coordination could extend beyond LC, as infraslow neuronal activities were recently identified in dorsomedial medulla that projects to LC.^63^ Furthermore, the direct demonstration of LC’s role in HR variability will renew interest in the varied autonomic and central manifestations of arousal-like events during NREMS.^44^

We find that mammalian sleep harnesses on wake promotion to enable sensory vigilance. This insight requires a renewal of current models of sleep-wake control in which reciprocal and exclusive antagonism is prevalent between sleep- and wake-promoting brain areas, including the LC.^8^ Still, the origins of how LC becomes periodically activated and overcomes this antagonism remain to be addressed. The LC is engaged in multiple and recurrent input-output loops across forebrain and hindbrain that could give rise to slow rhythms.^48^ Other possibilities to test are fluctuations in chemosensory signals that regulate LC^64, 65^ and that could in turn result from alterations in brain fluid transport during NREMS.^66^ Based on such insights, it could be interesting to probe whether the enhanced appearance of cyclic alternating patterns in cases of sleep-related breathing disorders such as sleep apnea^67^ can be controlled with noradrenergic antagonists. Furthermore, there is strong evidence that LC and sleep disruptions could be linked in post-traumatic stress disorders,^68^ in neurodegenerative diseases such as Alzheimer’s and Parkinson’s disease^45^ and in insomnia.^3^ We are now able to concretize questions into possible noradrenergic origins of a large variety of primary and secondary sleep disorders, in which hyperarousals, autonomic arousals and movement-related arousals prominently feature.^3, 44–46^

## Author contributions

Conceptualization, A.L., A.O.F. R.C., L.F.; Methodology, A.L., A.O.F.; Software, A.O.F., R.C; Validation, A.L, A.O.F., R.C, L.F.; Formal Analysis, A.O.F.; Investigation, A.O.F., R.C, G.V., A.G.G., G.K.; Resources, A.L., A.O.F.; Data curation, A.O.F., L.F.; Writing – Original Draft, A.L., A.O.F.; Writing – Review & Editing, L.F., R.C., G.V., A.G.G., G.K., A.O.F, A.L.; Visualization, A.O.F., L.F.; Supervision, A.L.; Project Administration, A.L.; Funding Acquisition, A.L., A.O.F.

## Acknowledgements

We thank Paul Steffan and Dr. David McCormick for providing us with the original DBH-Cre breeders. We greatly appreciate the time-efficient support provided to us from Drs. Yulong Li and Jessie Feng regarding the NE sniffers. Particular thanks go to the Animal caretaking team headed by Michelle Blom, and to Titouan Tromme who took so good care of our animal lines. Expert veterinary advice was given by Drs. Delphine Perret and Laure Sériot. We thank Christiane Devenoges for excellent support in histological analysis. We appreciate the helpful exchange regarding optogenetic LC manipulation and NE sniffer experimentation with Drs. Simone Astori, Antoine Adamantidis, Oxana Eschenko, Jessie Feng, Paul Franken, Celia Kjaerby, Maiken Nedergaard, Ernesto Durán, Andrea Volterra. We are indebted to Francesca Siclari and Christoph Michel for insightful discussions on the human noradrenergic system. Simone Astori, Francesca Siclari, and Freddy Weber provided valuable input on preliminary versions of the manuscript. All lab members provided constructive input to this study throughout the experimental period and to preliminary versions of the manuscript. This study was funded by The Swiss National Science Foundation (n° 310030-184759 to AL), Etat de Vaud and a FBM UNIL PhD Fellowship to AOF.

## STAR Methods

### RESOURCE AVAILABILITY

#### Lead contact

For information and request for resources should be directed to the lead contact, Anita Lüthi

#### Data and code availability

All data included in this publication will be stored on one of the servers of the University of Lausanne and will be made available once the study is published upon lead contact request. Customized MATLAB scripts for data acquisition and analysis are available from the corresponding author upon reasonable request.

#### Materials availability

The study did not produce any new materials or reagents.

### EXPERIMENTAL MODEL AND SUBJECT DETAILS

#### Subjects

Mice from the C57BL/6J line and from the B6.FVB(Cg)-Tg(Dbh-cre)KH212Gsat/Mmucd (MMRRC Stock#036778-UCD) line, referred to here as DBH-Cre line, were bred on a C57BL/6J background and housed in a humidity- and temperature-controlled animal house with a 12 h / 12 h light-dark cycle (lights on at 9 am). Food and water were available ad libitum throughout all the experimental procedures. For viral injections, 2- to 7-week-old mice of either sex were transferred to a P2 safety level housing room with identical conditions 1 d prior to injection. For in vivo experimentation, animals were transferred to the recording room 3 d after viral injection and left to recover for at least 1 week prior to the implantation surgery, after which they were singly housed in standard-sized cages. The grids on top of the cage were removed and replaced by 30 cm-high Plexiglass walls. Fresh food was regularly placed on the litter and the water bottle inserted through a hole in the cage wall. Objects (tissues, paper rolls, ping-pong balls) were given to play. For in vitro experimentation, animals were transferred 3 d after viral injection to a housing room with identical conditions and were used 3 – 6 weeks after injection. In total, 12 male C57BL/6J mice were used for intracranial pharmacological experiments, 19 male C57BL/6J mice for cardiac pharmacology experiments and 6 male C57BL/6J mice for the fiber photometry experiments. From the DBH-Cre line, 21 (11 males and 10 females) heterozygous Cre +/− animals were used for optogenetic experiments, and 14 (2 males and 12 females) for in vitro experiments. All experiments were conducted in accordance with the Swiss National Institutional Guidelines on Animal Experimentation and were approved by the Swiss Cantonal Veterinary Office Committee for Animal Experimentation.

### METHOD DETAILS

#### Viral injections

##### Optogenetics in vivo and in vitro

Animals were anaesthetized with ketamine (83 mg kg^−1^)/xylazine (3.5 mg kg^−1^), kept on a thermal blanket to maintain body temperature around 37 °C, and injected i.p. with carprofen (5 mg kg^−1^) for analgesia. Mice were then head-fixed on a stereotactic frame equipped with a head adaptor for young animals (Stoelting 51925). The scalp was disinfected, injected with a mix of lidocaine (6 mg kg^−1^)/bupivacaine (2.5 mg kg^−1^) for local anesthesia and opened with scissors exposing the desired region of the skull. For the injections, we used a thin glass pipette (5-000-1001-X, Drummond Scientific) pulled on a vertical puller (Narishige), initially filled with mineral oil, and backfilled with the virus-containing solution just prior to injection. Injections took place at an injection rate of 100 – 200 nl min^−1^. For optogenetic stimulation experiments, 2 animals were injected with a ssAAV5/2-hEF1α-dlox-hChR2(H134R)_mCherry(rev)-dlox-WPRE-hGHp(A) (titer: 9.1×10^12^ vg / ml, 0.8 – 1 µL; Zurich VVF) virus bilaterally in a region close to the LC. The stereotaxic coordinates were (relative to Bregma, given in mm here and throughout the rest of the STAR Methods): lateral (L) ±1.28; antero-posterior (AP) −5.45, depth (D) −3.65), as done previously.^9^ The remaining 9 animals were injected bilaterally with the same virus (0.3 – 0.6 µL) directly into the LC (L ±1.05; AP −5.45; D −3.06). The two viral injections yielded comparable results and data were pooled. For optogenetic inhibition, all animals were injected bilaterally into the LC (0.2 – 0.35 µL) with either pAAV5-CAG-FLEX-rc[Jaws-KGC-GFP-ER2] (7×10^12^ vg / ml; n = 2, Addgene), AAV8-hSyn-FLEX-Jaws-KGC-GFP-ER2 (3.2×10^12^ vg / ml; n = 3, UNC Vector Core) or ssAAV-5/2-hSyn1-dlox-Jaws_KGC_EGFP_ERES(rev)-dlox-WPRE-bGHp(A)-SV40p(A) (titer: 6.4×10^12^ vg / ml; n = 5, VVF Zürich).

##### Fiber photometry

For the assessment of NE dynamics in the thalamus, AAV viruses (ssAAV9/2-hSyn1-GRAB_NE1h-WPRE-hGHp(A), titer: 7.2×10^12^ vg / ml, or ssAAV9/2-hSyn1-GRAB_NE1m-WPRE-hGHp(A), titer: 5.5×10^12^ vg / ml, both from VVF Zürich) containing the plasmid encoding a NE sensor (pAAV-hSyn-GRAB_NE1h, Addgene Plasmid #123309, or pAAV-hSyn-GRAB_NE1m, Addgene Plasmid #123308, respectively)^28^ were injected into the thalamus (500 nl; L 2.0; AP −1.6; D −3.0). After the injections, the incision was sutured, and the area disinfected. Animals were carefully monitored and returned to the home cage once awake and moving around. Recovery time after injections took place for a minimum of 1 week before the next surgeries. Paracetamol was given in the water for the 4 postoperative days at a concentration of 2 mg ml^−1^.

#### Other surgical procedures

For in vivo EEG/EMG combined with local field potential (LFP) recordings, electrode implantation was as previously described.^18, 21, 27^ In short, animals were anesthetized with isoflurane (1.5 – 2.5 %) in a mixture of O_2_ and N_2_O. After analgesia (i.p. carprofen 5 mg kg^−1^) and disinfection, animals were fixed in a Kopf stereotax and injected into the scalp with a mix of lidocaine (6 mg kg^−1^)/bupivacaine (2.5 mg kg^−1^) and a piece of the scalp was removed after 3 – 5 min, the skull exposed and the bone scratched to improve adhesion of the head implant. Then, we drilled small craniotomies (0.3 – 0.5 mm) over left frontal and parietal bones and positioned two conventional gold-coated wire electrodes in contact with the dura mater for EEG recordings. On the contralateral (right) side, a high-impedance tungsten LFP microelectrode (10–12 MΩ, 75 µm shaft diameter, FHC) was implanted in the primary somatosensory cortex (L 3; AP −0.7; D - 0.85). Additionally, as a neutral reference, a silver wire (Harvard Apparatus) was inserted into the occipital bone over the cerebellum and two gold pellets were inserted into the neck muscles for EMG recordings. All electrodes were fixed using Loctite Schnellkleber 401 glue and soldered to a multisite connector (Barrettes Connectors 1.27 mm, male connectors, Conrad).

For intracranial injection of noradrenergic antagonists, we additionally made a craniotomy over the thalamus (L 2; AP −1.60) and covered it with a silicone-based sealant (Kwik-Cast Silicone Sealant, WPI). Additionally, we glued and cemented a light-weight metal head-post (Bourgeois Mécanique SAS, Lyon, France) onto the midline skull to perform painless head-fixation during injection of noradrenergic antagonists.

For optogenetic experiments, DBH-Cre animals were implanted with custom-made optic fibers.^69^ A multimode fiber (225 μm outer diameter, Thorlabs, BFL37-2000/FT200EMT) was inserted and glued (heat-curable epoxy, Precision Fiber Products, ET-353ND-16OZ) to a multimode ceramic zirconia ferrule (Precision Fiber Products, MM-FER2007C-2300). The penetrating end was cut at the desired length with a carbide-tip fiber optic scribe (Precision Fiber Products, M1-46124). The outside end was then polished using fiber-polishing films (Thorlabs). For optogenetic stimulation of the LC cell bodies (n = 11 animals) a single 3-mm fiber stub was implanted directly over the LC (L 1.0; AP −5.4; D −2.3). Out of the 11 animals, 6 animals were also implanted with a 3 mm-optic fiber stub over the somatosensory thalamus (L 2.0; AP - 1.7; D −2.5). For the 5 additional animals, we implanted a custom-made optrode in S1 built with a high-impedance fine tungsten LFP microelectrodes (10 – 12 MΩ, 75 µm shaft diameter, FHC) glued to the stub of a 2 mm-optic fiber at a distance of 800 – 1,200 µm. The optrode was then inserted into S1 (L 3.0; AP - 0.7; D 0.8). For optogenetic inhibition of the LC bodies (n = 10), bilateral optic fibers were implanted at a 20° lateral angle targeting the LC (L ±1.84; AP −5.4; D −2.47). To establish the final coordinates of the optic fibers, pupil diameter changes were monitored in a subgroup of 5 animals while lowering the optic fiber and applying light stimuli (Figure S4; 10 – 30 pulses at 10 Hz). A custom-made software developed in Matlab was used for image acquisition (See in vivo data analysis).

For fiber photometry experiments, in addition to the recording electrodes, we implanted C57BL/6J animals (n = 6) with a premade 400 µm-thick optic fiber coupled to a cannula (MFC_400/430-0.66_3.5mm_ZF1 25(G)_FLT, Doris Lenses) over the dorsal and reticular thalamus (L 1.8; AP −1.7; D 2.5) at a speed of 1 mm min^−1^.

Finally, a dental cement structure was built to fix the implant in place. After disinfection with iodine-based cream, animals were returned to their home cage and kept in careful monitoring. Animals were provided with paracetamol (2 mg mL^−1^) in the drinking water for at least 4 days after the procedure.

#### In vivo electrophysiological recordings

Once recovered from the surgery, animals were habituated to the cabling for 5 – 7 days, followed by a baseline recording to ascertain the quality of the signals. We acquired the EEG, EMG and LFP signals at a 1 kHz sampling frequency using an Intan digital RHD2132 amplifier board and a RHD2000 USB Interface board (Intan Technologies) connected via SPI cables (RHD recording system, Intan Technologies). Homemade adapters containing an Omnetics - A79022-001 connector (Omnetics Connector Corp.) linked to a female Barrettes Connector (Conrad) were used as an intermediate between the head implant of the animal and the headstage. We acquired the data with Matlab using the RHD2000 Matlab toolbox and a customized software in the same environment.^21, 27^

#### Procedures for intracranial pharmacology

We gently and gradually habituated mice to being head-fixed by increasing the amount of time spent in head fixation daily from 5 min to 45 min over a period of 4 – 5 days. The rest of the time, the animals spent being tethered to the recording system in their home cage. On the first experimental day, we removed the silicone cover of the craniotomy in head-fixed conditions and positioned a glass pipette (5-000-1001-X, Drummond Scientific, pulled on a vertical Narishige PP-830 puller, tip size of 15 – 25 µm) over the craniotomy and waited for 30 min while gently touching the side of the craniotomy to simulate an injection. Then, we covered the craniotomy again with the silicone-based sealant and returned the animals to the home cage for an 8 h-baseline polysomnographic recording. The next day, we removed the silicone again and injected 150 nL of noradrenergic antagonists or ACSF at two different depths within the thalamus (D: −3.2 and −2.8 mm). For the experimental group, we infused a mixture of 0.1 mM prazosin hydrochloride (prazosin) and 5 mM (S)-(-)-atenolol (atenolol), diluted in ACSF together with a red fluorescent dye (5 mM Alexa 594) for later confirmation of the injection site. For the control group, we injected ACSF together with Alexa 594. Per animal, only one injection was done (either blockers or ACSF) and the animal sacrificed after completion of the recording.

#### Procedures for in vivo optogenetics

All optogenetic manipulation took place during the first 20 min of each hour between ZT1 and ZT9. A custom-made close-loop detection of NREMS^27^ was used for state specificity. In short, NREMS was detected whenever the delta (1 – 4 Hz) to theta (5 – 10 Hz) power ratio derived from the differential frontal-parietal EEG channels crossed a threshold for 2 out of 5 s and the EMG absolute values went below a threshold during at least 3 s. This lead to reliable stimulation during NREMS (Figure 2), with the exception of a few brief interruptions that occurred during artefacts (e.g. muscle twitches). Stimulation sessions took place in the first 20 min of each hour during 8 h of the light phase (ZT1-9), with light or sham (light source turned off) stimulation alternating over successive recording days. Optogenetic stimulation of the LC cell bodies was carried out using a PlexBright Optogenetic Stimulation System (Plexon) coupled to a PlexBright Table-top blue LED Module (Wavelength 465 nm) at 1 Hz. Stimulation of LC terminals in the thalamus or cortex was delivered at 2 Hz (Figure 3). Optogenetic inhibition of LC cell bodies was performed using a continuous stimulation with a PlexBright Table-top orange LED Module (Wavelength 620 nm) (Figure 2). Optogenetic stimulation (1 Hz) or inhibition (continuous) was also carried out specifically when sigma power declined or rose. This was achieved via a machine-learning-based closed-loop procedure^27^ built with a multilayer perceptron model neural network of 10 neurons in the hidden layer and 3 output neurons (for rising or decreasing sigma power during NREM or to no stimulation, in the case of epochs outside this sleep state). The network was fed with the last 200 s of a 9^th^-order polynomial fit of the sigma activity (10 – 15 Hz) calculated for each s. The neural network was then trained, validated, and tested using the sleep scoring from 13 C57BL/6J animals (642,000 epochs) that were otherwise not included in this study. Online, the same data stretches obtained from the mouse in recording were used.

For each animal, multiple recording sessions took place with a random allocation of the stimulation protocol: i.e. optogenetic stimulation during NREMS in the LC bodies, its terminals (thalamus for 6 animals or cortex 5 animals), or stimulation of the LC bodies during spindle-enriched or -poor substates (in a subgroup of 9 mice). Similarly, for optogenetic inhibition of LC bodies, random allocation of inhibition protocols took place during NREMS (10 animals) or NREMS substates (10 animals).

#### Procedures for in vivo fiber photometry

After the recovery (> 7 d) and habituation to the cabling procedure (> 4 d), we performed two recordings per animal with at least one day between sessions. All recordings were limited to the first 3 – 4 h from ZT1 to minimize possible photobleaching. For fluorescent measurements, we used a pulse-width-modulated sinusoidal signal of 400 Hz using a Raspberry Pi3 (Raspberry Pi Foundation) to modulate a LEDD_2 driver (Doric Lenses Inc.) connected to a blue LED (CLED 465 nm; Doric Lenses Inc.). The power of the driver was set to 200 mA. The blue LED was coupled to a fluorescence MiniCube (iFMC4_IE(400-410)_E(E460-490)_F(500-550)_S, Doric Lenses) that redirected the light to the animal via a low autofluorescence 400-µm-thick fiberoptic patchcord (MFP_400/430/1100-0.57_1m_FMC-ZF1.25_LAF, Doric Lenses Inc.). The cord was connected to the Optic fiberoptic Cannula (MFC_400/430-0.57_3mm_ZF1.25(G)_FLT) implanted in the head of the mouse. A photodetector integrated into the MiniCube head turned the emitted light from the fluorescent NE sensor into a current signal that was fed into an analog signal of the Intan RHD2132 amplifier board. To ensure that data collected were within the dynamic range of the biosensors, awake animals were exposed to the experimenter’s hand held within the cage for 1 min moving gently but without touching the animal (See Figure S5).

#### In vitro electrophysiological recordings

Thalamic brain slice recordings were performed as previously described in detail.^21, 69^ Briefly, 3 – 6 weeks after viral injection, DBH-Cre mice aged 8 – 16 weeks were subjected to isoflurane anesthesia, after which they were decapitated, brains extracted and quickly immersed in ice-cold oxygenated sucrose solution (which contained in mM): NaCl 66, KCl 2.5, NaH_2_PO_4_ 1.25, NaHCO_3_ 26, D-saccharose 105, D-glucose 27, L(+)-ascorbic acid 1.7, CaCl_2_ 0.5 and MgCl_2_ 7), using a sliding vibratome (Histocom). Brains were trimmed at the level of the brainstem, glued on the trimmed surface on an ice-cold metal blade and apposed to a supporting agar block on their ventral side. Acute 300-μm-thick coronal brain slices were prepared in the same ice-cold oxygenated sucrose solution and kept for 30 min in a recovery solution at 35 °C (in mM: NaCl 131, KCl 2.5, NaH_2_PO_4_ 1.25, NaHCO_3_ 26, D-glucose 20, L(+)-ascorbic acid 1.7, CaCl_2_ 2, MgCl_2_ 1.2, *myo*-inositol 3, pyruvate 2) before being transferred to room temperature for at least 30 min. All recordings were done at room temperature.

Recording glass pipettes were pulled from borosilicate glass (TW150F-4) (WPI) with a DMZ horizontal puller (Zeitz Instr.) to a final resistance of 2 – 4 MΩ. Pipettes were filled with a K^+^-based intracellular solution that contained in mM: KGluconate 140, Hepes 10, KCl 10, EGTA 0.1, phosphocreatine 10, Mg-ATP 4, Na-GTP 0.4, pH 7.3, 290–305 mOsm. Slices were placed in the recording chamber of an upright microscope (Olympus BX50WI) and continuously superfused with oxygenated ACSF containing in mM: NaCl 131, KCl 2.5, NaH_2_PO_4_ 1.25, NaHCO_3_ 26, D-glucose 20, L(+)-ascorbic acid 1.7, CaCl_2_ 2 and MgCl_2_ 1.2. Cells were visualized with differential interference contrast optics and 10X and 40X immersion objectives, and their location within the thalamic ventroposterial medial nucleus or within the somatosensory reticular thalamus could be verified based on previous studies in the lab.^21, 69^ Infrared images were acquired with an iXon Camera X2481 (Andor). Prior to recording, pipette offset was zeroed, and the stability of the offset verified by monitoring pipette potential in the bath for 10 min. Drifts were < 0.5 mV / 10 min. Signals were amplified using a Multiclamp 700B amplifier, digitized via a Digidata1322A and sampled at 10 kHz with Clampex10.2 (Molecular Devices). Immediately after gaining whole-cell access, cellular membrane potential and access resistance were measured. Cells included had a resting membrane potential < −55 mV and access resistances < 15 MΩ. The cell types were identified based on their rebound bursting properties (See Figure S6). Whole-field blue LED (Cairn Res) stimulation (455 or 470 nm, duration: 0.1 – 1 ms, maximal light intensity 0.16 and 0.75 mW/mm^2^ for the two LEDs, respectively). Per slice, only one cell was recorded and exposed to light stimulation. For characterization of LC fiber-evoked membrane depolarizations, cells were held between −65 to −70 mV and exposed to 1 Hz, 3 Hz or 10 Hz stimulation (4 pulses each). Stimulation at different frequencies were applied in random order, with each frequency used maximally twice to avoid run-down of the evoked response. When light-induced depolarizations did not return to the original membrane potential, they were not included in the analysis. For the study of LC-dependent effects on prolonged afterdepolarizations, thalamocortical cells were first injected with series of repetitive negative current injections (100 – 300 pA, 20 pulses, each 120 ms) known to evoke rebound low-threshold Ca^2+^ bursts. Such protocols have been used previously to characterize the cell-intrinsic mechanisms accompanying sleep-spindle-related arrival of barrages of inhibitory synaptic potentials.^30^ Following 1 – 2 such repetitive current injections (each followed by 3 – 5 min of recovery time), the current injections were preceded by LC fiber stimulation (10 Hz, 4 pulses) by 5 s, such that the maximum of the LC-evoked membrane depolarization coincided with the end of the negative current injections. For characterization of LC fiber-evoked membrane currents, cells were held in voltage-clamp at −70 mV. Baseline light-evoked currents were evoked maximally 1 – 2 times, followed by bath application of cesium chloride or noradrenergic antagonists ((S)-(-)-atenolol (Abcam) for thalamocortical cells or prazosin hydrochloride (Abcam) for thalamic reticular cells) for 5 – 10 min before the next optogenetic stimulation. The in vitro data were manually analyzed using Clampfit v2.2 and as illustrated in Figure S6.

#### Pharmacological manipulation of heart rate

After the recovery period of the electrode implantation (> 7 d), mice were habituated to the recording conditions for one week. Mice were injected intraperitoneally with NaCl, (S)-(-)-atenolol (1 mg kg^−1^) (Abcam), a sympathetic antagonist or methylatropine bromide (10 mg kg^−1^) (Sigma-Aldrich), a parasympathetic antagonist, both known to poorly permeate the blood-brain barrier.^31, 32^ Injections were done at 9 am and followed by polysomnographic recording for 100 min. Two recording sessions per drug took place in an intercalated manner. Experimenters were blind to the drug injected.

#### Histology

After all recording sessions were completed, animals were injected intraperitoneally with a lethal dose of pentobarbital. For animals implanted with electrodes for LFP recording, the position of the electrode was marked via electro-coagulation (50 µA, 8 – 10 s) of the region. Subsequently, ∼45 mL of paraformaldehyde (PFA) 4% were perfused intracardially at a rate of ∼2.5 mL min^−1^. Brains were post-fixed for at least 24 h in PFA 4% cooled to 4 °C. Brains were then sliced in 100 µm-thick sections with a vibratome (Microtome Leica VT1000 S; speed: 0.25 – 0.5 mm s^−1^ and knife sectioning frequency: 65 Hz) or a freezing microtome (Microm). Brain sections were directly mounted on slides or kept in well plates filled with 0.1 M PB for later processing. Then, we confirmed the position of LFP electrodes and optic fibers and the fluorescent expression of the injected viruses or local pharmacology injections with a Nikon SMZ25 Stereomicroscope equipped with a Nikon DS-Ri2 16 Mpx color camera. When needed, higher magnification images were acquired using an Axiovision Imager Z1 (Zeiss) microscope equipped with an AxioCam MRc5 camera (objectives used EC-Plan Neofluar 2.5x/0.075 ∞/0.17, 5x/0.16 ∞/0.17, 10x/0.3 ∞/− or 20x/0.5 ∞/0.17).

### QUANTIFICATION AND STATISTICAL ANALYSIS

#### In vivo data analysis

##### Scoring of vigilance states

We detected sleep and wake episodes following previous standard procedures in a manner blinded to the treatment.^21, 27^ For this purpose, we used a custom-made software developed in Matlab (MathWorks) that allows semiautomatic scoring of sleep stages. Shortly, we defined three distinct stages as follows: wakefulness, periods containing large muscle tonus or phasic activity in the EMG signal, together with low-voltage EEG exhibiting fast oscillatory components. NREMS was defined as periods containing low EMG activity together with high amplitude EEG activity showing slow oscillatory components such as slow oscillations (< 1.5 Hz), delta (1.5 – 4 Hz) or sleep spindles (10 – 15 Hz). REMS episodes were defined as periods with low EMG activity with prominent Theta (5 – 10 Hz) activity in the EEG. Microarousals were defined as short (< 12 s) periods of wakefulness contained between the epochs of the same sleep stage. For the intracranial pharmacology experiments, analysis was done for the first 2 h of recording and comparisons were made in a paired manner between the baseline and drug conditions. For all optogenetic experiments, scored data for the first 20 min of each hour (during which light stimulation was done) were compared with the same periods in sham conditions (*ceteris paribus* with the LED turned off). For the fiber photometry experiments, analysis included the complete 3 – 4 h of recordings. For the pharmacological manipulation of the HR, analysis took place for the first 100 min after the i.p. injections.

##### Analysis of sigma and delta dynamics

Dynamics of sigma (10 – 15 Hz) and delta (1.5 – 4 Hz) activities were quantified from the S1 LFP signal using a wavelet transform with a Mother Gabor-Morlet wavelet with 4 cycles of standard deviation for the Gaussian envelope (see Figures 1A and 1C). The frequency dimensions were then collapsed to the two frequency bands of interest, the 10 – 15 Hz sigma band and the 1.5 – 4 Hz delta band. The mean signals were then resampled at 10 Hz and filtered using a 100^th^ order filter with a 0.025 Hz cutoff frequency for further analysis. For NREMS bouts of ≥ 96 s, a Fast Fourier transform was calculated (Figure 1B) to measure the strength of the 0.02 Hz oscillatory patterns defined here as the area underneath the Fourier transform from 0.01 – 0.04 Hz, subtracting the mean activity between 0.08 to 0.12 Hz (as depicted in Figure 1B). For the example depicted in Figure 1B, the sigma and delta grand averages were calculated from the long bouts (> 96 s) of NREMS of baseline recordings contained within ZT1-7 (local pharmacology experiment) or ZT1-9 (optogenetic experiments).

##### Sleep spindle detection, phase coupling analysis and feature extraction

Sleep spindles were detected from the S1 LFP signal for all the experiments. Spindle detection was done using a previously described algorithm^21^ that is illustrated in Figure S1. Briefly, we filtered (FIR filter of order 2000) the raw S1 LFP signal in the sigma band (9 – 16 Hz). Then, we squared the signal and applied a threshold of 1.5 the standard deviation above the mean values in NREMS. We then detected all the peaks crossing this threshold and marked as a putative spindle all events containing at least 3 cycles. The starting and ending point of the events were extended to the closest cycle at 0 crossing before and after the threshold, respectively. Events separated by < 50 ms were merged as a single event. For display purposes, we positioned a black dot for each individual spindle event at the center of the spindle in time and a random jittered vertical position.

After spindle detection, we extracted the following features: *Amplitude,* the maximum value of the absolute filtered signal within the event. *Frequency,* mean intra-peak frequency within the detected spindle event. *Number of cycles* in the spindle. *Duration,* timespan between the beginning and end of the event.

For the analysis of the phase coupling of the spindle events to the sigma activity we first centered the sigma activity at zero by subtracting the mean of the sigma activity in NREMS. Then we constructed a distribution using the phase of the sigma dynamics (calculated as described in *Analysis of sigma and delta dynamics*) at the center of each spindle event (half point between the beginning and the end of the spindle). By using the CircStat toolbox for Matlab (MathWorks),^70^ we then confirmed the non-uniformity of the distribution by using the Rayleigh test.

##### Detection of infraslow cycles

We detected individual cycles of sigma within NREMS using a custom-made Matlab routine. We used the sigma dynamics as described in “*Analysis of sigma and delta dynamics”* and eliminated the regions containing artifacts. Then, we identified the peaks and troughs in the signal with a minimum distance of 25 and 20 s, respectively. Finally, we arranged the positions of successive troughs and kept the starting and ending point for each individual cycle. Next, the marked locations were used to normalize the time in 1000 points for each individual cycle and to interpolate the sigma activity to generate a mean dynamics normalized in time. The same positions were used to normalize the dynamics of NE-related fluorescent signals from the fiber photometry measurements. Only those cycles that within NREMS periods were included in the mean.

##### Pupil diameter measurements

To standardize the correct location of the optic fibers in the LC cell bodies stimulation or inhibition, we performed pupil diameter measurements in a subset of animals. In short, we set a Basler GigE infrared camera (Basler acA800-510 um, SVGA, 1/3.6’’, 510 fps, USB3 Vision) close to one eye of the animal and used a custom-made infrared LED-based lantern directed to the recorded eye to increase the contrast between the pupil and the surrounded area. We built a custom-made software in Matlab (MathWorks) for online or offline pupil detection (using the videos recorded from online trials). First, the user manually selects both the area of interest for the analysis and the initial location of the pupil. Then, for each frame (recorded at 10 fps), a binary image was created using an Otsu’s method adaptive threshold with the function `imbinarize` from Matlab (MathWorks). The threshold was set manually to adapt to the conditions of the image and the angles of the infrared light source and the camera. The size of the binary object closest to the marked pupil was then measured. The dynamics of the pupil diameter was then tracked and z-scored for comparison (See Figure S4).

##### Fiber Photometry

Changes in bioluminescence were recorded via a photodetector connected to an analog channel in the Intan RHD2000 USB Interface board (Intan Technologies) as described before. The recorded signal fluctuated between 0 – 3.3 V with peaks at 400 Hz as the sinusoidal waveform created to modulate the excitation of the biosensors. The fluorescent dynamics signal was then created using an RMS envelope of 1 s. The changes in biofluorescence ΔF * F^−1^ were computed by dividing the enveloped signal by its fitted exponential decrease calculated from the dynamics at NREMS. We computed the relative fluorescence across different sleep states using the z-scored data per recording session; the values presented in Figure 5B show the mean values of multiple sessions per animal for quiet wakefulness (QW), NREM (NR) and REM (R) sleep. In every case, only epochs flanked by epochs of the same vigilance state were included. *Cross correlation analysis:* To study the similarity of the dynamics between the sigma activity and the NE changes or changes in HR, we performed cross correlation analysis between these two pairs of signals in Matlab (MathWorks). For each long bout (≥ 96 s) of NREMS within the time of analysis (see *Scoring of vigilance states*), we z-scored each signal individually and normalized it to the length of the bout. Then, the cross-correlation was calculated using the function `xcorr` of Matlab and normalized to the length of the bout. Mean cross correlation values were computed for each animal and across animals. Mean correlation coefficient (r) was computed between −5 to 5 s lags.

##### Heart rate analysis

Changes in HR were computed as previously described.^18^ Shortly, EMG signal was filtered using a Chebyshev type 2 high-pass filter. We then differentiated (using the function `diff` from Matlab) and squared the signal to highlight the R peaks. The resulting signal was z-scored to normalize across animals and recording sessions. Then, the R peaks were identified using the Matlab function `findpeaks` with a heuristically found threshold of 0.3 and an interpeak distance of at least 0.08 ms. Peaks with a z-score higher than 10 were considered as artifacts and eliminated. Finally, the HR signal was constructed by measuring the inter-peak time distance and divided by 60 (1 min). An interpolation of the values was performed using the function `interp1` of Matlab and resampled at 10 Hz.

#### Statistical analysis

For the statistical tests, we used R statistical language version 3.6.1. and Matlab (MathWorks). First, we tested for normality of the datasets using the Shapiro-Wilk normality test. For comparisons of two parametric datasets, we used a paired Student’s t test and the equivalent Wilcoxon signed rank test for non-parametric datasets. For comparisons on multiple (> 2) groups of data (as in the case of the amplitude, onset latency and recovery time in the in-vitro experiments), a one-way ANOVA or a Kruskal-Wallis test was used for parametric and non-parametric datasets, respectively. In case of no significance, no further *post-hoc* analysis was performed. In all figures, grey lines denote paired datasets from two conditions (e.g. baseline, opto). Mean values are given by large horizontal lines, error bars indicate standard errors of the mean.

### KEY RESOURCES TABLE

| REAGENT or RESOURCE | SOURCE | IDENTIFIER |
| --- | --- | --- |
| **Bacterial and virus strains** |  |  |
| ssAAV5/2-hEF1α-dlox-hChr2(H134R)_mCherry(rev)-dlox-WPRE-hGHp(A) | VVF Zurich | v80-5 |
| pAAV5-CAG-FLEX-rc[Jaws-KGC-GFP-ER2] | Addgene | 84445-AAV5 |
| AAV8-hSyn-FLEX-Jaws-KGC-GFP-ER2 | UNC Vector Core | N/A |
| ssAAV-5/2-hSyn1-dlox-Jaws_KGC_EGFP_ERES(rev)-dlox-WPRE-bGHp(A)-SV40p(A) | VVF Zurich | v508-5 |
| ssAAV9/2-hSyn1-GRAB_NEh-WPRE-hGHp(A) | VVF Zurich | v472-9 |
| ssAAV9/2-hSyn1-GRAB_NEm-WPRE-hGHp(A) | VVF Zurich | v471-9 |
| **Chemicals, peptides, and recombinant proteins** |  |  |
| (S)-(-)-atenolol | Sigma-Aldrich | A7655 |
| Prazosin hydrochloride | Sigma-Aldrich | P7791 |
| Methylatropine bromide | Sigma-Aldrich | M1300000 |
| Cesium chloride | Sigma-Aldrich | C3032-25G |
| **Recombinant DNA** |  |  |
| pAAV-hSyn-GRAB_NE1h | Addgene | Plasmid #123309 |
| pAAV-hSyn-GRAB_NE1m | Addgene | Plasmid #123308 |
| **Deposited data** |  |  |
| Raw and analyzed data | This paper |  |
| MATLAB code supporting HD and MWM analyses | This paper | available from lead contact on request |
| **Experimental models: Organisms/strains** |  |  |
| M. musculus C57BL/6J | Jackson Laboratory | Jax: 000664 |
| M. musculus B6.FVB(Cg)-Tg(Dbh-cre)KH212Gsat/Mmucd | MMRRC, University of California, Davis | MMRRC_036778-UCD |
| **Software and algorithms** |  |  |
| NIS-Elements 4.5 | Nikon | N/A |
| Zen lite 2012 | Zeiss | N/A |
| Adobe Photoshop 2020 | Adobe Creative Cloud | N/A |
| Adobe Illustrator 2020, 2021 | Adobe Creative Cloud | N/A |
| Clampex10.2 | Molecular Devices | N/A |
| Clampfit v2.2 | Molecular Devices | N/A |
| R 3.5.1 | R Core Team | N/A |
| MATLAB 2015b, 2018a | MathWorks | N/A |
| Excel | Microsoft | N/A |

## Supplemental Information

**This PDF file includes:**

**Figure S1.**
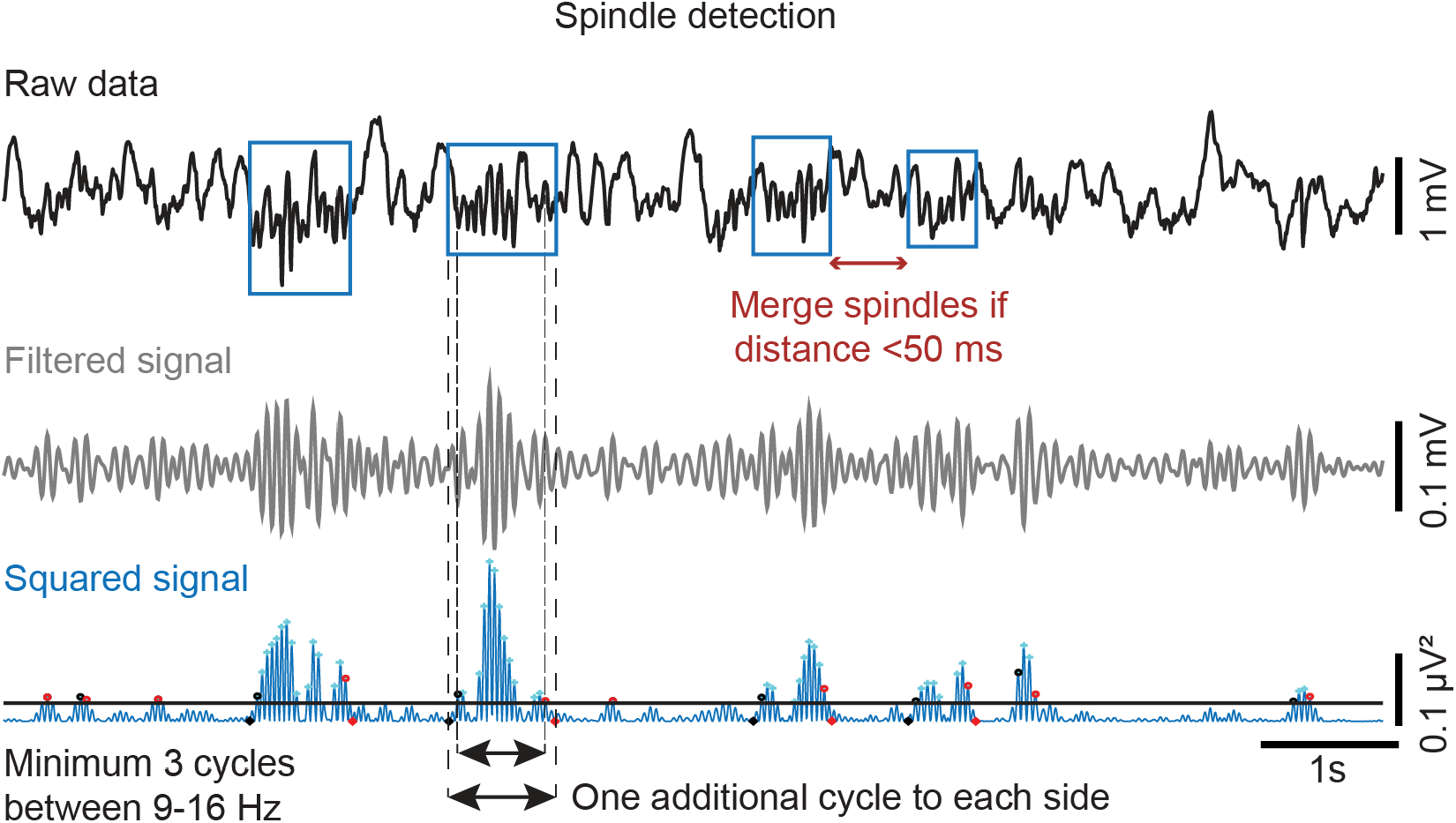
Automated detection of individual spindles. Raw data from S1 LFP recordings (top), filtered (middle) and squared signals (bottom) were used to extract individual spindles according to the criteria given next to the traces. The method was identical to the one published recently [S1].

**Figure S2.**
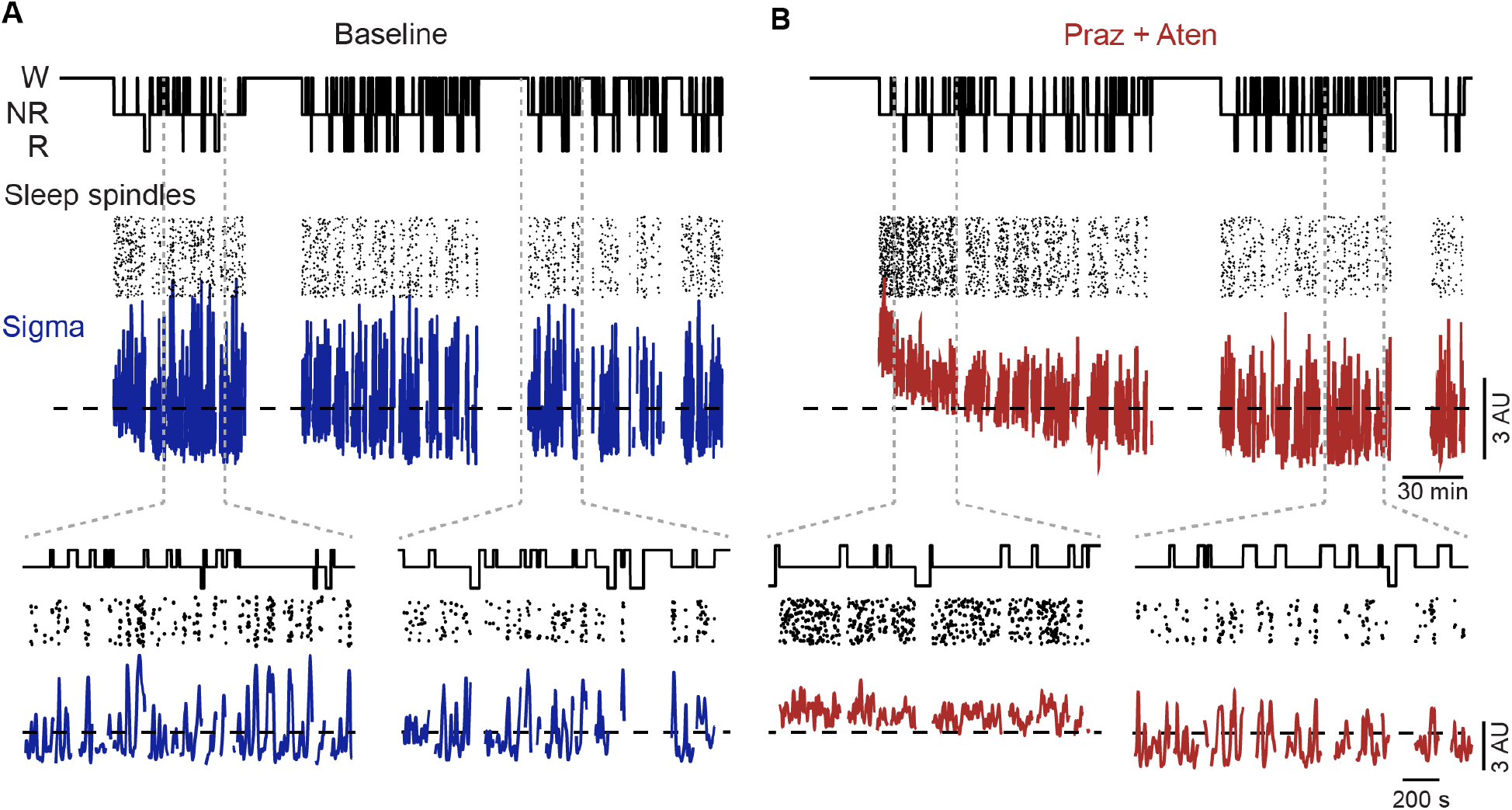
Time course of sigma power and sleep spindle density during pharmacological antagonism of noradrenergic signaling in thalamus. (A) Animal in baseline conditions, showing, from top to bottom, hypnogram, individual sleep spindles, and sigma power dynamics, with expanded portions shown below. (B), Same Animal, injected with Prazosin (Praz) and Atenolol (Aten). Note slow recovery of the pharmacological suppression of sigma power fluctuations and sleep spindle clustering. Every animal was injected once with either antagonists or ACSF.

**Figure S3.**
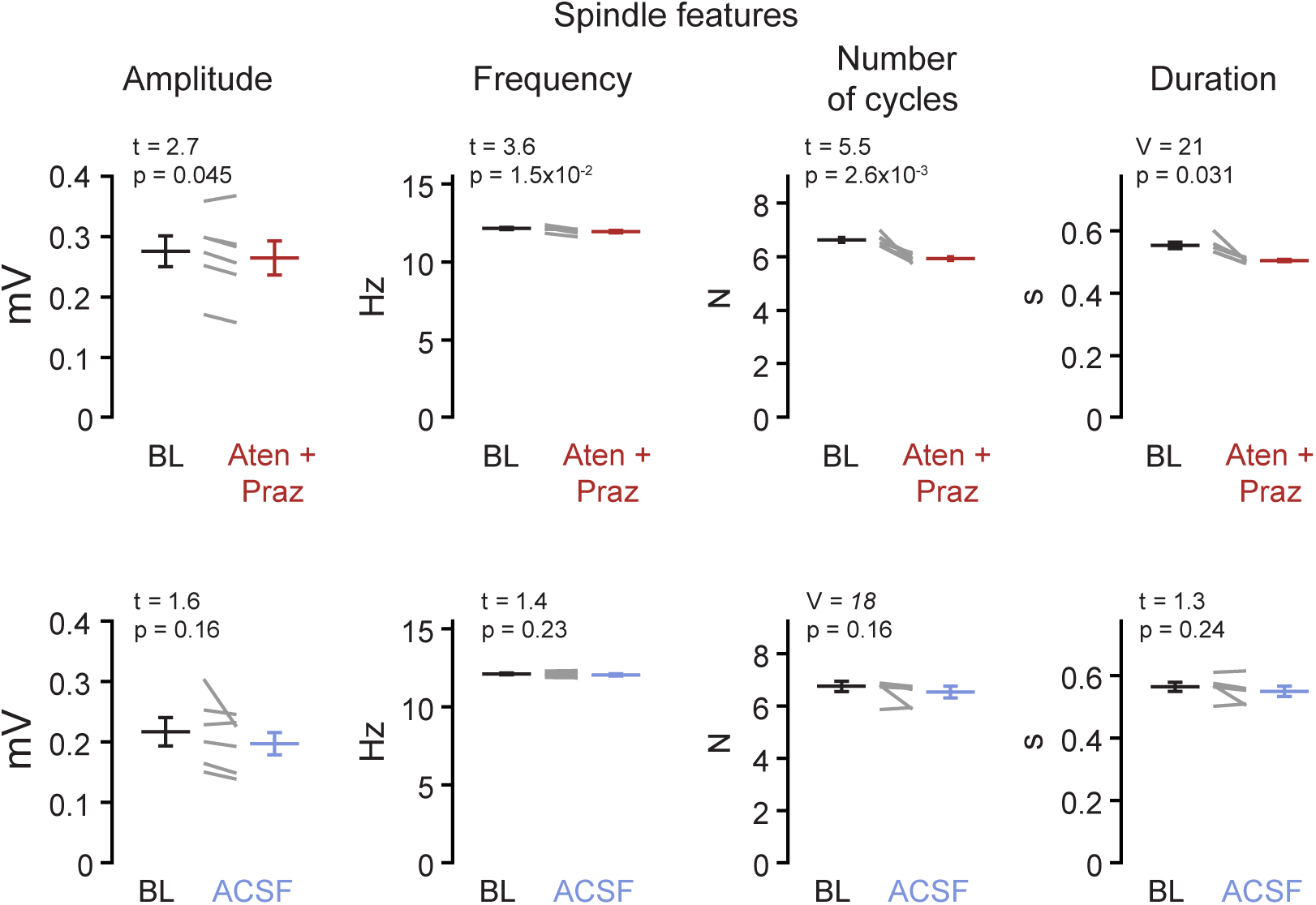
Quantification of properties of individual sleep spindles in animals used for the pharmacology experiments. Top; Data from the 6 animals injected with noradrenergic antagonists Atenolol (Aten) and Prazosin (Praz), Bottom; Data from the 6 animals injected with ACSF. From left to right: Peak amplitude, Intra-spindle frequency, Number of spindle cycles, and total spindle duration, analyzed as illustrated in Figure S1 and according to ref. [S1]. V, t and p in this figure were obtained from Wilcoxon signed rank or paired t-tests.

**Figure S4.**
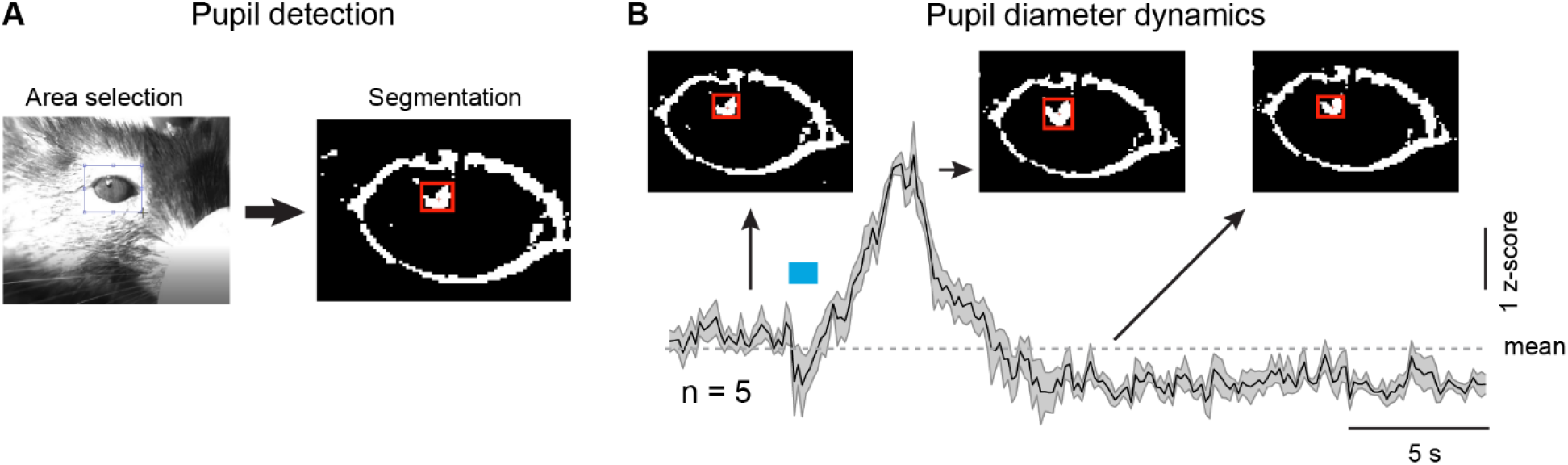
LC stimulation-induced pupil diameter changes. (A) Representative image taken from the mouse eye during surgical implantation of optic fiber, showing area selection (dashed square) and corresponding image segmentation with pixels selected for measurement (red square). (B) Quantification of time course of pupil diameter changes in z-score, with representative segmented images in insets. Blue bar denotes light application. These recordings were made in a subgroup of 5 animals to optimize the position of the optic fiber for subsequent LC optogenetic manipulation during NREMS.

**Figure S5.**
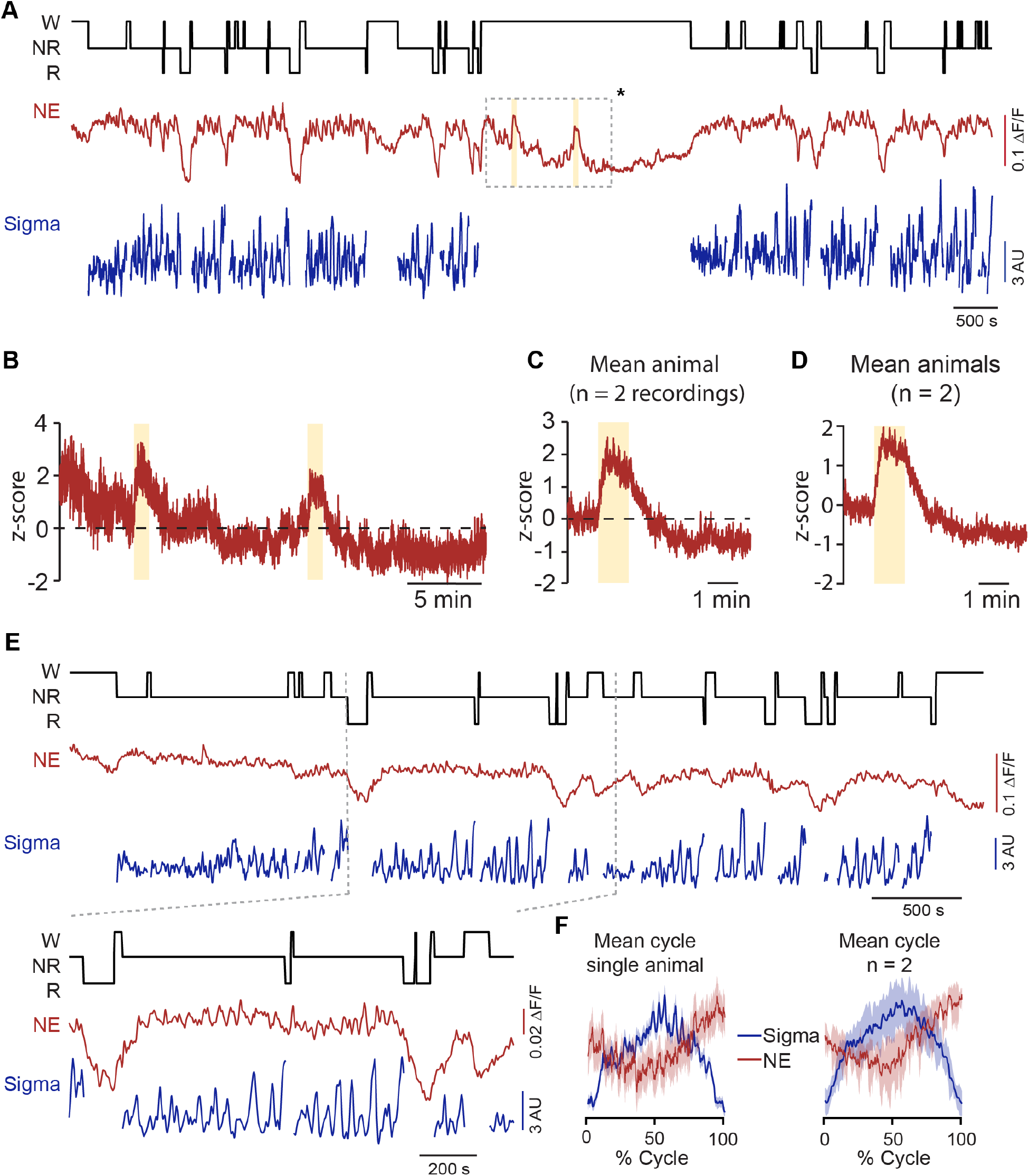
Further results ascertaining the suitability of the NE biosensors for tracking free NE dynamics during the sleep-wake cycle. (A) Representative recording showing (from top to bottom) hypnogram, relative fluorescence derived from the NE biosensor GRAB_NE1h_, and sigma power dynamics. Areas labeled with yellow vertical line represent 1-min bouts during which the experimenter’s hand was gently moved inside the cage of the animal. Note the increase of NE when this perturbation took place. (B) Expanded z-scored portion from the marked (*) region in (A). (C) Mean NE increase produced by hand insertion in two recordings from the same animal. (D) Mean NE increase across two animals. (E) Representative traces as in (A), showing the relative fluorescence measured with the medium-affinity NE biosensor GRAB_NE1m_. Expanded portion shown below for the marked area. (F) Left, overlay of sigma power dynamics across all infraslow cycles detected from trough to trough, with corresponding NE fiber photometry signal for one mouse measured with the medium affinity biosensor GRAB_NE1m_. Right, Mean traces across two animals with the same biosensor. Note the similar time course compared to the ones shown for the high-affinity sensor in Figure 5.

**Figure S6.**
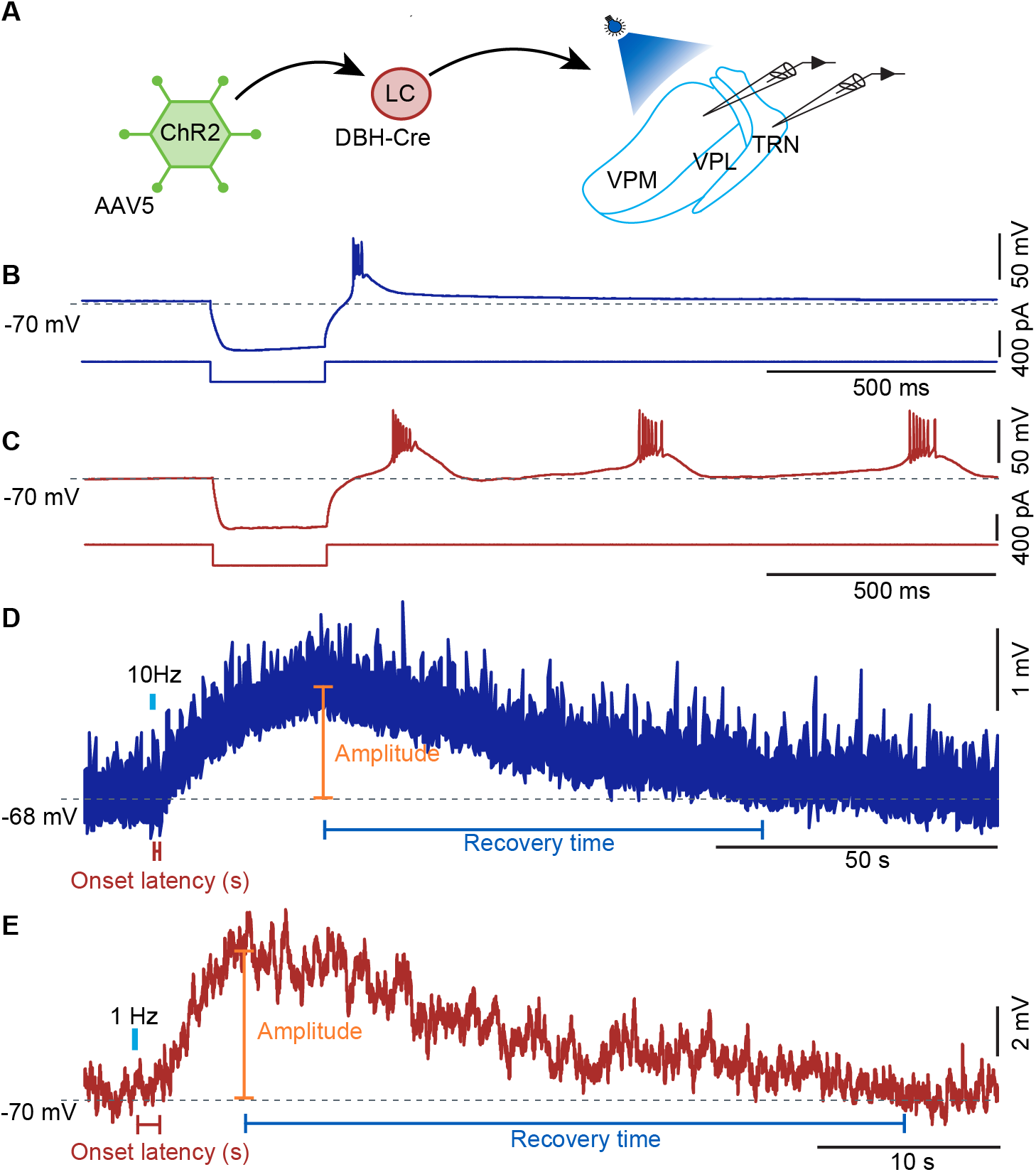
Details for the in vitro patch-clamp recordings and analysis. (A) Schematic showing the experimental design. VPM, ventroposterior medial nucleus, VPL, ventroposterior lateral nucleus, TRN, thalamic reticular nucleus. Recordings were performed in either VPM or TRN while light was shone to activate afferent LC fibers. (B) Rebound bursting characteristic for a thalamocortical cell recorded in VPM and evoked through brief negative current injection. (C) Repetitive rebound bursting characteristic for a TRN cell recorded in the somatosensory sector and evoked through brief negative current injection. (D) Enlarged light-evoked response of a thalamocortical neuron, with indications on how quantification of onset latency, amplitude and recovery time were done. (E) As (D), for a light-evoked response in TRN. Dashed lines denote membrane potential value indicated on the left.

## References

1. Van Someren, E.J., Cirelli, C., Dijk, D.J., Van Cauter, E., Schwartz, S., and Chee, M.W. (2015). Disrupted sleep: from molecules to cognition. J Neurosci 35, 13889–13895.

2. Mascetti, G.G. (2020). Adaptation and survival: hypotheses about the neural mechanisms of unihemispheric sleep. Laterality, 1–23.

3. Van Someren, E.J.W. (2020). Brain mechanisms of insomnia: new perspectives on causes and consequences. Physiol Rev, in press.

4. Siclari, F., Valli, K., and Arnulf, I. (2020). Dreams and nightmares in healthy adults and in patients with sleep and neurological disorders. Lancet Neurol 19, 849–859.

5. Mander, B.A., Winer, J.R., and Walker, M.P. (2017). Sleep and human aging. Neuron 94, 19–36.

6. Saper, C.B., Fuller, P.M., Pedersen, N.P., Lu, J., and Scammell, T.E. (2010). Sleep state switching. Neuron 68, 1023–1042.

7. Aston-Jones, G., and Bloom, F.E. (1981). Activity of norepinephrine-containing locus coeruleus neurons in behaving rats anticipates fluctuations in the sleep-waking cycle. J Neurosci 1, 876–886.

8. Saper, C.B., and Fuller, P.M. (2017). Wake-sleep circuitry: an overview. Curr Opin Neurobiol 44, 186–192.

9. Carter, M.E., Yizhar, O., Chikahisa, S., Nguyen, H., Adamantidis, A., Nishino, S., Deisseroth, K., and de Lecea, L. (2010). Tuning arousal with optogenetic modulation of locus coeruleus neurons. Nat Neurosci 13, 1526–1533.

10. Poe, G.R., Foote, S., Eschenko, O., Johansen, J.P., Bouret, S., Aston-Jones, G., Harley, C.W., Manahan-Vaughan, D., Weinshenker, D., Valentino, R., et al. (2020). Locus coeruleus: a new look at the blue spot. Nat Rev Neurosci 21, 644–659.

11. Eschenko, O., and Sara, S.J. (2008). Learning-dependent, transient increase of activity in noradrenergic neurons of locus coeruleus during slow wave sleep in the rat: brain stem-cortex interplay for memory consolidation? Cereb Cortex 18, 2596–2603.

12. Swift, K.M., Gross, B.A., Frazer, M.A., Bauer, D.S., Clark, K.J.D., Vazey, E.M., Aston-Jones, G., Li, Y., Pickering, A.E., Sara, S.J., et al. (2018). Abnormal locus coeruleus sleep activity alters sleep signatures of memory consolidation and impairs place cell stability and spatial memory. Curr Biol 28, 3599–3609 e3594.

13. Kjaerby, C., Andersen, M., Hauglund, N., Ding, F., Wang, W., Xu, Q., Deng, S., Kang, N., Peng, S., Sun, Q., et al. (2020). Dynamic fluctuations of the locus coeruleus-norepinephrine system underlie sleep state transitions. Biorxiv, 2020.2009.2001.274977.

14. Dang-Vu, T.T., Schabus, M., Desseilles, M., Albouy, G., Boly, M., Darsaud, A., Gais, S., Rauchs, G., Sterpenich, V., Vandewalle, G., et al. (2008). Spontaneous neural activity during human slow wave sleep. Proc Natl Acad Sci U S A 105, 15160–15165.

15. Gais, S., Rasch, B., Dahmen, J.C., Sara, S., and Born, J. (2011). The memory function of noradrenergic activity in non-REM sleep. J Cogn Neurosci 23, 2582–2592.

16. Hayat, H., Regev, N., Matosevich, N., Sales, A., Paredes-Rodriguez, E., Krom, A.J., Bergman, L., Li, Y., Lavigne, M., Kremer, E.J., et al. (2020). Locus coeruleus norepinephrine activity mediates sensory-evoked awakenings from sleep. Sci Adv 6, eaaz4232.

17. Zerbi, V., Floriou-Servou, A., Markicevic, M., Vermeiren, Y., Sturman, O., Privitera, M., von Ziegler, L., Ferrari, K.D., Weber, B., De Deyn, P.P., et al. (2019). Rapid reconfiguration of the functional connectome after chemogenetic *locus coeruleus* activation. Neuron 103, 702–718 e705.

18. Lecci, S., Fernandez, L.M.J., Weber, F.D., Cardis, R., Chatton, J.-Y., Born, J., and Lüthi, A. (2017). Coordinated infra-slow neural and cardiac oscillations mark fragility and offline periods in mammalian sleep. Sci. Adv. 3, e1602026.

19. Fernandez, L.M.J., and Lüthi, A. (2020). Sleep spindles: mechanisms and functions. Physiol Rev 100, 805–868.

20. Andrillon, T., and Kouider, S. (2020). The vigilant sleeper: neural mechanisms of sensory (de)coupling during sleep. Curr Opin Physiol 15, 47–59.

21. Fernandez, L.M., Vantomme, G., Osorio-Forero, A., Cardis, R., Béard, E., and Lüthi, A. (2018). Thalamic reticular control of local sleep in sensory cortex. Elife 7, e39111.

22. Lázár, Z.I., Dijk, D.J., and Lázár, A.S. (2019). Infraslow oscillations in human sleep spindle activity. J Neurosci Methods 316, 22–34.

23. Csernai, M., Borbély, S., Kocsis, K., Burka, D., Fekete, Z., Balogh, V., Káli, S., Emri, Z., and Barthó, P. (2019). Dynamics of sleep oscillations is coupled to brain temperature on multiple scales. J Physiol 597, 4069–4086.

24. Eschenko, O., Magri, C., Panzeri, S., and Sara, S.J. (2012). Noradrenergic neurons of the locus coeruleus are phase locked to cortical up-down states during sleep. Cereb Cortex 22, 426–435.

25. Wyatt, R.J., Chase, T.N., Kupfer, D.J., Scott, J., and Snyder, F. (1971). Brain catecholamines and human sleep. Nature 233, 63–65.

26. Agster, K.L., Mejias-Aponte, C.A., Clark, B.D., and Waterhouse, B.D. (2013). Evidence for a regional specificity in the density and distribution of noradrenergic varicosities in rat cortex. J Comp Neurol 521, 2195–2207.

27. Cardis, R., Lecci, S., Fernandez, L.M.J., Osorio-Forero, A., Chu Sin Chung, P., Fulda, S., Decosterd, I., and Lüthi, A. (2021). Local cortical arousals and heightened (somato)sensory arousability during non-REM sleep of mice in chronic pain. Biorxiv, 2021.2001.2004.425347.

28. Feng, J., Zhang, C., Lischinsky, J.E., Jing, M., Zhou, J., Wang, H., Zhang, Y., Dong, A., Wu, Z., Wu, H., et al. (2019). A genetically encoded fluorescent sensor for rapid and specific *in vivo* detection of norepinephrine. Neuron 102, 745–761 e748.

29. Lee, K.H., and McCormick, D.A. (1996). Abolition of spindle oscillations by serotonin and norepinephrine in the ferret lateral geniculate and perigeniculate nuclei *in vitro*. Neuron 17, 309–321.

30. Lüthi, A., and McCormick, D.A. (1999). Modulation of a pacemaker current through Ca^2+^-induced stimulation of cAMP production. Nat Neurosci 2, 634–641.

31. Smith, R.D., Grzelak, M.E., and Coffin, V.L. (1994). Methylatropine blocks the central effects of cholinergic antagonists. Behav Pharmacol 5, 167–175.

32. Neil-Dwyer, G., Bartlett, J., McAinsh, J., and Cruickshank, J.M. (1981). β-adrenoceptor blockers and the blood-brain barrier. Br J Clin Pharmacol 11, 549–553.

33. Mather, M., Joo Yoo, H., Clewett, D.V., Lee, T.H., Greening, S.G., Ponzio, A., Min, J., and Thayer, J.F. (2017). Higher locus coeruleus MRI contrast is associated with lower parasympathetic influence over heart rate variability. Neuroimage 150, 329–335.

34. Wang, X., Pinol, R.A., Byrne, P., and Mendelowitz, D. (2014). Optogenetic stimulation of locus ceruleus neurons augments inhibitory transmission to parasympathetic cardiac vagal neurons via activation of brainstem α1 and β1 receptors. J Neurosci 34, 6182–6189.

35. Smeets, W.J., and Gonzalez, A. (2000). Catecholamine systems in the brain of vertebrates: new perspectives through a comparative approach. Brain Res Brain Res Rev 33, 308–379.

36. Noronha-de-Souza, C.R., Bicego, K.C., Michel, G., Glass, M.L., Branco, L.G., and Gargaglioni, L.H. (2006). Locus coeruleus is a central chemoreceptive site in toads. Am J Physiol Regul Integr Comp Physiol 291, R997–1006.

37. Totah, N.K.B., Logothetis, N.K., and Eschenko, O. (2019). Noradrenergic ensemble-based modulation of cognition over multiple timescales. Brain Res 1709, 50–66.

38. Bouret, S., and Sara, S.J. (2005). Network reset: a simplified overarching theory of locus coeruleus noradrenaline function. Trends Neurosci 28, 574–582.

39. Yüzgeç, O., Prsa, M., Zimmermann, R., and Huber, D. (2018). Pupil size coupling to cortical states protects the stability of deep sleep via parasympathetic modulation. Curr Biol 28, 392–400 e393.

40. Watson, B.O. (2018). Cognitive and physiologic impacts of the infraslow oscillation. Front Syst Neurosci 12, 44.

41. Libourel, P.A., Barrillot, B., Arthaud, S., Massot, B., Morel, A.L., Beuf, O., Herrel, A., and Luppi, P.H. (2018). Partial homologies between sleep states in lizards, mammals, and birds suggest a complex evolution of sleep states in amniotes. PLoS Biol 16, e2005982.

42. Antony, J.W., Piloto, L., Wang, M., Pacheco, P., Norman, K.A., and Paller, K.A. (2018). Sleep spindle refractoriness segregates periods of memory reactivation. Curr Biol 28, 1736–1743.

43. Klinzing, J.G., Niethard, N., and Born, J. (2019). Mechanisms of systems memory consolidation during sleep. Nat Neurosci 22, 1598–1610.

44. Parrino, L., Halasz, P., Tassinari, C.A., and Terzano, M.G. (2006). CAP, epilepsy and motor events during sleep: the unifying role of arousal. Sleep Med Rev 10, 267–285.

45. Doppler, C.E.J., Smit, J.A.M., Hommelsen, M., Seger, A., Horsager, J., Kinnerup, M.B., Hansen, A.K., Fedorova, T.D., Knudsen, K., Otto, M., et al. (2021). Microsleep disturbances are associated with noradrenergic dysfunction in Parkinson’s disease. Sleep, in press.

46. Ferri, R., Koo, B.B., Picchietti, D.L., and Fulda, S. (2017). Periodic leg movements during sleep: phenotype, neurophysiology, and clinical significance. Sleep Med 31, 29–38.

47. Totah, N.K., Neves, R.M., Panzeri, S., Logothetis, N.K., and Eschenko, O. (2018). The locus coeruleus is a complex and differentiated neuromodulatory system. Neuron 99, 1055–1068 e1056.

48. Schwarz, L.A., and Luo, L. (2015). Organization of the locus coeruleus-norepinephrine system. Curr Biol 25, R1051–R1056.

49. Niethard, N., Ngo, H.V., Ehrlich, I., and Born, J. (2018). Cortical circuit activity underlying sleep slow oscillations and spindles. Proc Natl Acad Sci U S A 115, E9220–E9229.

50. Seibt, J., Richard, C.J., Sigl-Glöckner, J., Takahashi, N., Kaplan, D.I., Doron, G., de Limoges, D., Bocklisch, C., and Larkum, M.E. (2017). Cortical dendritic activity correlates with spindle-rich oscillations during sleep in rodents. Nat Commun 8, 684.

51. Niethard, N., Brodt, S., and Born, J. (2021). Cell-type specific dynamics of calcium activity in cortical circuits over the course of slow wave sleep and rapid eye movement sleep. J Neurosci, in press.

52. Jones, B.E. (2020). Arousal and sleep circuits. Neuropsychopharmacology 45, 6–20.

53. el Mansari, M., Sakai, K., and Jouvet, M. (1989). Unitary characteristics of presumptive cholinergic tegmental neurons during the sleep-waking cycle in freely moving cats. Exp Brain Res 76, 519–529.

54. Kayama, Y., Ohta, M., and Jodo, E. (1992). Firing of ‘possibly’ cholinergic neurons in the rat laterodorsal tegmental nucleus during sleep and wakefulness. Brain Res 569, 210–220.

55. Durán, E., Oyanedel, C.N., Niethard, N., Inostroza, M., and Born, J. (2018). Sleep stage dynamics in neocortex and hippocampus. Sleep 41, zsy060.

56. Takahashi, K., Kayama, Y., Lin, J.S., and Sakai, K. (2010). Locus coeruleus neuronal activity during the sleep-waking cycle in mice. Neuroscience 169, 1115–1126.

57. Gaspar, P., Berger, B., Febvret, A., Vigny, A., and Henry, J.P. (1989). Catecholamine innervation of the human cerebral cortex as revealed by comparative immunohistochemistry of tyrosine hydroxylase and dopamine-beta-hydroxylase. J Comp Neurol 279, 249–271.

58. Hermans, E.J., van Marle, H.J., Ossewaarde, L., Henckens, M.J., Qin, S., van Kesteren, M.T., Schoots, V.C., Cousijn, H., Rijpkema, M., Oostenveld, R., et al. (2011). Stress-related noradrenergic activity prompts large-scale neural network reconfiguration. Science 334, 1151–1153.

59. Lüthi, A., and McCormick, D.A. (1998). Periodicity of thalamic synchronized oscillations: the role of Ca^2+^-mediated upregulation of I_h_. Neuron 20, 553–563.

60. Hughes, S.W., Lörincz, M.L., Parri, H.R., and Crunelli, V. (2011). Infraslow (<0.1 Hz) oscillations in thalamic relay nuclei: basic mechanisms and significance to health and disease states. Prog Brain Res 193, 145–162.

61. Yang, B., Sanches-Padilla, J., Kondapalli, J., Morison, S.L., Delpire, E., Awatramani, R., and Surmeier, D.J. (2021). Locus coeruleus anchors a trisynaptic circuit controlling fear-induced suppression of feeding. Neuron 109, 1–16.

62. Nothias, F., Onteniente, B., Roudier, F., and Peschanksi, M. (1988). Immunocytochemical study of serotoninergic and noradrenergic innervation of the ventrobasal complex of the rat thalamus. Neurosci Lett 95, 59–63.

63. Stucynski, J.A., Schott, A.L., Baik, J., Chung, S., and Weber, F. (2021). Regulation of REM sleep by inhibitory neurons in the dorsomedial medulla. Biorxiv 2020.2011.2030.405530.

64. Li, A., and Nattie, E. (2006). Catecholamine neurones in rats modulate sleep, breathing, central chemoreception and breathing variability. J Physiol 570, 385–396.

65. Madan, V., and Jha, S.K. (2012). A moderate increase of physiological CO_2_ in a critical range during stable NREM sleep episode: a potential gateway to REM sleep. Front Neurol 3, 19.

66. Fultz, N.E., Bonmassar, G., Setsompop, K., Stickgold, R.A., Rosen, B.R., Polimeni, J.R., and Lewis, L.D. (2019). Coupled electrophysiological, hemodynamic, and cerebrospinal fluid oscillations in human sleep. Science 366, 628–631.

67. Parrino, L., Ferri, R., Bruni, O., and Terzano, M.G. (2012). Cyclic alternating pattern (CAP): the marker of sleep instability. Sleep Med Rev 16, 27–45.

68. Reist, C., Streja, E., Tang, C.C., Shapiro, B., Mintz, J., and Hollifield, M. (2020). Prazosin for treatment of post-traumatic stress disorder: a systematic review and meta-analysis. CNS Spectr, 1–7.

69. Vantomme, G., Rovó, Z., Cardis, R., Béard, E., Katsioudi, G., Guadagno, A., Perrenoud, V., Fernandez, L.M.J., and Lüthi, A. (2020). A thalamic reticular circuit for head direction cell tuning and spatial navigation. Cell Rep 31, 107747.

70. Berens, P. (2009). CircStat: A MATLAB toolbox for circular statistics. J Stat Software 31, 1–21.

## Supplemental References

S1. L. M. Fernandez, G. Vantomme, A. Osorio-Forero, R. Cardis, E. Béard, A. Lüthi, (2018). Thalamic reticular control of local sleep in sensory cortex. Elife 7, e39111.

